# Individual olfactory channels shape distinct parameters of sleep architecture

**DOI:** 10.1101/2025.09.11.675654

**Authors:** Oliver M. Cook, Benjamin T. Pursley, Garrett Pauls, Stephen Roeder, Isaiah A. Aites, Kaylynn E. Coates, Andrew M. Dacks

## Abstract

Across the animal kingdom, olfactory dysfunction and anosmia have been associated with disruptions in sleep. In the fruit fly *Drosophila melanogaster*, various studies have demonstrated that broadly inhibiting olfactory receptor neurons (ORNs) similarly disrupts sleep/wake cycles, suggesting that baseline ORN signaling is an integral component of olfactory modulation of sleep. However, due to the diversity of ORNs and combinatorial nature of olfactory processing, many of the cellular and molecular mechanisms by which ORNs modulate sleep remain unclear. In this study, we addressed this gap of knowledge by characterizing the contributions of different ensembles of ORNs, individual ORN types, and a known modulator of ORNs on baseline sleep architecture. We find that the activity of distinct ORN types are important for day and nighttime sleep and heterogeneously shape parameters of sleep architecture. Importantly, the effects of ORN signaling on sleep are adjusted across mating status, suggesting that distinct ORN types are recruited within the context of sleep depending on the demands of the animal. Furthermore, the effects of ORN signaling on sleep are in part shaped by heterogeneous serotonin (5-HT) receptor expression. Together, this work identifies cellular and molecular pathways bridging olfaction and sleep, and helps establish a circuit model that can be used to further characterize the behavioral consequences of sensory dysfunction.

## Introduction

As animals navigate their environment they actively rely on their sense of smell to locate food, avoid predators/parasites/disease, and mate with conspecifics^1–7^. However, there is now emerging evidence that olfactory processing is also important during quiescent or sleeping periods, ensuring that animals can adjust their arousal states in the presence of threatening or beneficial stimuli^8–11^. Odor stimulations can either improve or disrupt sleep depending on the odor type and the context in which they are presented^12–19^. This suggests that the activity of distinct olfactory pathways may flexibly modulate baseline sleep architecture. Indeed, olfactory dysfunction and anosmia are correlated with disruptions in sleep/wake cycles and may even influence the onset and progression of sleep disorders in patients with Parkinson’s disease^20–22^. However, despite this apparent relationship between olfaction and sleep, it has been difficult to ascertain the extent to which olfactory dysfunction causally affects sleep health. Namely, the presence of comorbid conditions in Parkinson’s generally overshadows any direct contributions of olfactory signaling on sleep. Additionally, given the combinatorial nature of olfactory processing^23,24^, identifying specific sleep-modulating olfactory pathways can be difficult. For example, since hyposmic/anosmic individuals do not always experience the same olfactory deficits ^25,26^, sleep architecture may be shaped in a variety of ways depending on the pathways dysregulated across individuals. Therefore, in order to gain a more mechanistic understanding of olfactory modulation of sleep, it is important to leverage genetically tractable animal models in which identified olfactory circuitry can be manipulated.

In mice, the removal of the olfactory bulb (the primary olfactory center in mammals) disrupts distinct phases of sleep architecture^27^, suggesting that olfactory modulation of sleep may first arise in the primary processing stages. Similarly in the fruit fly *Drosophila melanogaster,* physical ablation or genetic silencing of olfactory receptor neurons (ORNs)^28–30^ (which transduce odorant cues from the environment to the brain^31^), as well as knockouts of genes related to ORN signaling^32^ each disrupt different parameters of sleep architecture, suggesting that ORNs are a key cell type involved in olfactory modulation of sleep. Thus, ORNs and their synaptic partners are an excellent model to explore the fundamental properties of olfactory modulation of sleep, especially given advancements in circuit tracing and functional characterization of olfactory cells^33–43^. With this model we can ask questions such as: Do specific ORN pathways modulate distinct parameters of sleep, or is there a generalized role distributed across all ORNs? Do the effects of ORN signaling on sleep vary across different internal and external contexts of the animal? What upstream signaling pathways modulate ORNs within the context of sleep? Addressing these questions would provide a mechanistic framework as to how olfactory signaling shapes sleep architecture.

The goal of this study was to explore the cellular and molecular mechanisms by which ORN signaling affects baseline sleep architecture. We address this question by characterizing the contributions of ensembles of ORNs, individual ORN types, and a known sleep modulator, serotonin. By using drivers of largely non-overlapping ORN types, we first determined that specific ORN types modulate sleep. Notably, distinct ORN types modulate different temporal phases and parameters of sleep architecture, rather than all ORNs having a uniform impact. Additionally, the contributions of some ORN classes varies across mating status, consistent with previous reports of context-dependent changes in ORN excitability. Finally, serotonin signaling to ORNs recapitulates many of the same responses, indicating that the influence of olfaction on sleep can be up- or downregulated depending on the ORN pathway being modulated. Together, this work reveals cellular and molecular mechanisms underlying the relationships between olfaction and sleep and provides a circuit model for future studies characterizing the impacts of sensory dysfunction on sleep architecture.

## Results

The olfactory system processes odorants through a combinatorial code^23^, such that odor detection and subsequent representation in the brain are largely dependent on the differential activation patterns of ORN types^33,44,45^. There are ∼60 ORN types in *Drosophila*^46^, each responsible for transducing different odor scenes to the brain, classified by the chemoreceptive tuning receptors (mostly “*Or*” or *Ir*”)^47–52^ and co-receptors (*Ir8a*, *Ir25a*, *Ir76b*, and *Orco*)^53–59^ that they express. Removal of the third antennal segment in which the majority of ORNs are housed, or thermogenetically silencing all *Orco+* ORNs at once, both reduce total sleep duration^30^. Considering the combinatorial nature of olfaction as it may relate to sleep, these findings suggest that either 1) *Orco*+ ORNs are the predominant types that modulate sleep and that individual ORN types have redundant functions, or 2) broadly downregulating ORN signaling masks the different contributions of each ORN type.

### Distinct ORN co-receptor populations influence different parameters of day and nighttime sleep architecture

Our first question was whether the signaling from ORNs that expressed co-receptors other than *Orco* modulated sleep (**Figure 1A**). To test this hypothesis, we first constitutively silenced largely non-overlapping populations of ORN types by driving the expression of botulinum neurotoxin (BoNT-C)^60^ in different ORN co-receptor GAL4 lines. We chose to investigate the effects of silencing ORNs within *Orco*, *Ir8a*, and *Ir25a*-GAL4 drivers because these are the most widely expressed co-receptor types and there is little overlap between their expression patterns. *Orco*-GAL4 labels ORNs innervating the majority of the anterior glomeruli in the primary olfactory center of flies (known as the antennal lobes (AL))^55^ (**Figure 1B**). *Ir25a*-GAL4 labels ORNs innervating ∼12 glomeruli, the majority of which are posterior and function in hygro/thermosensation^61–64^, and other sensory cells throughout the body^65^ (**Figure 1C**). *Ir8a*-GAL4 labels ORNs innervating ∼10-12 posterior glomeruli that instead respond to appetitive odors including apple cider vinegar, acids, ammonia and polyamines^61,66–68^ **(Figure 1D)**.

**Figure 1.**
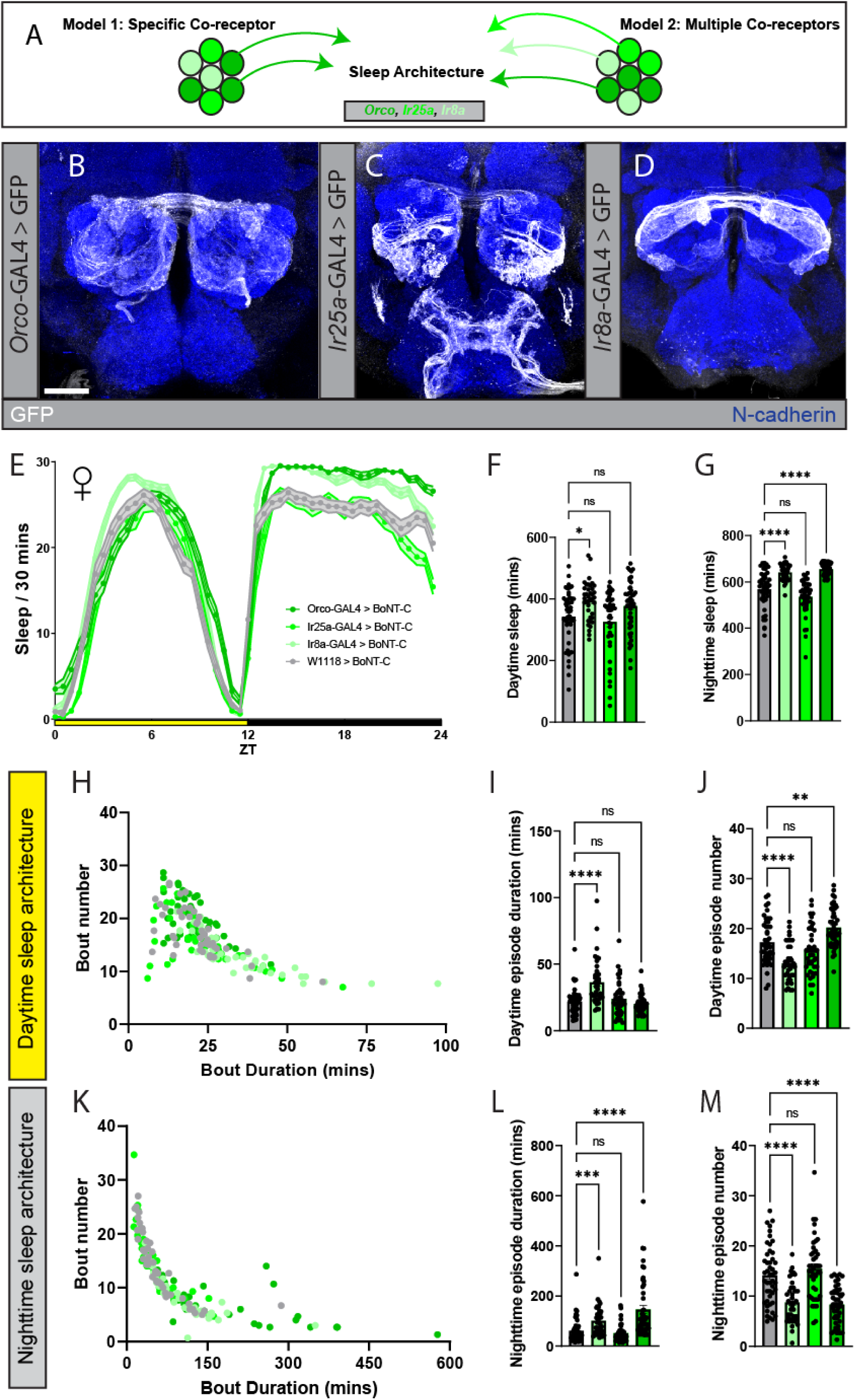
Neurotransmission of distinct ORN co-receptor populations modulates day or nighttime sleep architecture. **A**) Schematic of opposing models where either a distinct ORN co-receptor population, such as *Orco*+ ORNs, are the only types that modulate sleep vs one in which ORNs from various co-receptor populations influence sleep. **B-D)** Glomerular innervation patterns of ORNs within different ORN co-receptor populations: **B**) *Orco*-GAL4, **C**) *Ir25a*-GAL4, and **D**) *Ir8a*-GAL4. GFP signal is driven in the respective GAL4 driver (white), while the neuropil is labeled by N-cadherin (blue). Scale bar is 30µm. **E**) Sleep per 30 minutes of flies where different ORN co-receptor populations are silenced via BoNT-C. Yellow bar represents daytime (ZT 0-12), while black is nighttime (ZT 12-24). **F-G**) Average sleep duration of **F**) daytime and **G**) nighttime sleep phases. **H**) Scatter plot of the per-fly daytime sleep architecture (comparison of daytime sleep bout duration and number) across each tested genotype. **I**) Average daytime sleep bout duration. **J**) Average daytime sleep bout number. **K**) Scatter plot of the per-fly nighttime sleep architecture (comparison of nighttime sleep bout or duration and number). **L**) Average nighttime sleep bout duration. **M**) Average nighttime sleep bout number. In each behavior experiment, 3-7 day age mated female flies were tested, N=39-47 per genotype. Gray = BoNT-C controls, lightest green = *Ir8a*-GAL4 > BoNT-C, brighter green = *Ir25a*-GAL4 > BoNT-C, darkest green = *Orco*-GAL4 > BoNT-C. See methods for details on statistical analysis. *p<0.05, **p<0.01, ***p<0.001, ****p<0.0001.

Constitutively silencing different ORN co-receptor populations heterogeneously modulated sleep in mated females (**Figure 1E**), indicating that *Orco*+ ORNs are not the only ORN types that influence sleep. Silencing *Ir8a*+ ORNs increased both the total daytime and nighttime sleep duration, whereas silencing *Orco*+ ORNs only increased nighttime sleep and silencing *Ir25a+* ORNs had no effect on sleep duration (**Figure 1F-G**). Additionally, some of these changes in sleep duration were accompanied by deficits of locomotor activity during waking periods. We observed a general trend that as the number of ORN types expressed in the driver increased, the activity per waking minute decreased. Thus, silencing *Orco*+ ORNs produced a strong deficit in daytime and nighttime activity per waking minute (**Supplemental Figure 1I-J**). Together, these results reveal two important points. First, the impact of olfaction on sleep is due to the activity of ORNs that respond to specific odors, rather than generalized olfactory sensitivity. Furthermore, signaling from distinct populations of ORN types have different influences on day and nighttime sleep, suggesting that specific odor channels make unique contributions to sleep architecture.

Sleep duration is shaped by several factors including the probability of sleep initiation and bout duration once sleep is entered^69^. Since silencing *Ir8a*+ and *Orco*+ ORNs both increased sleep duration, albeit at different times of day, we next questioned whether ORNs uniformly influence sleep architecture. We therefore assessed the impacts of silencing different ORN co-receptor populations on metrics of sleep architecture (sleep episode duration and number of sleep episodes), sleep depth (pwake) and sleep drive (pdoze)^70^ separately for daytime and nighttime sleep as each period is regulated by distinct circuitry^71^. Not surprisingly, silencing *Ir8a+* and *Orco+* ORNs affected different aspects of daytime sleep architecture. Silencing *Ir8a*+ ORNs consolidated daytime sleep architecture by increasing the daytime sleep episode duration and decreasing the sleep episode number (**Figure 1H-J)**, whereas *Orco*+ ORNs instead increased daytime sleep episode number independent of episode duration (**Figure 1H-J**). This suggests that silencing *Ir8a+* ORNs consolidated sleep, whereas silencing Orco+ ORNs increased sleep initiation (**Figure 1F**). Furthermore, silencing *Ir8a*+ ORNs decreased the probability to transition out of an active state in the daytime, decreasing daytime pwake (**Supplemental Figure 1A, 1C-D**), while silencing *Orco*+ ORNs increased the probability of transitioning to an inactive state in the daytime, increasing daytime pdoze (**Supplemental Figure 1B-C, 1E**). Overall, this indicated that silencing selective ORN populations modulates different parameters of sleep architecture with *Ir8a*+ ORNs affecting sleep consolidation and *Orco*+ ORNs affecting the probability of entering a sleep bout.

Just as silencing *Ir8a*+ and *Orco*+ ORNs both increased nighttime sleep, we also observed similar effects on several aspects of sleep architecture, suggesting that both manipulations play a role in consolidating sleep. Silencing each ORN population increased nighttime sleep bout duration, decreased sleep bout number (**Figure 1K-M**), and decreased nighttime pwake (**Supplemental Figure 1A, 1F-G**), although silencing both *Ir8a*+ and *Ir25a*+ ORNs decreased nighttime pdoze, while *Orco*+ ORNs had no effect (**Supplemental Figure 1B, 1F, 1H**). Altogether, this screen outlined two general principles of ORN sleep modulation: 1) Distinct ORN types impact sleep more than others. For instance, silencing *Ir8a*+ and *Orco*+ ORNs strongly influence sleep architecture, while *Ir25a*+ ORNs have little to no effect. 2) Distinct ORN types impact different temporal phases and parameters of sleep architecture. For instance, silencing *Ir8a*+ ORNs strongly consolidated daytime and nighttime sleep, whereas silencing *Orco*+ ORNs only consolidated nighttime.

### ORN sleep modulation changes across mating status

The internal state of an animal can alter the relevance of a given odor^72^. For example, many *Ir8a*+ ORNs are receptive to polyamines and food-related odors, which are more attractive to mated females than virgin females^73^. Hunger status and diet also alters the excitability of ORNs^74–76^, thus a variety of contexts could shape the overall contributions of ORN signaling on sleep. Since silencing the activity of distinct ORN types influenced sleep more than others, we hypothesized that changes in the internal state of the animal, and subsequently the value placed on specific odor cues, may alter the influence of ORNs on sleep in a cell type-specific manner. To test this hypothesis, we chose to repeat the ORN co-receptor BoNT-C screen within mated males and virgin females to determine if sex or mating status altered the contribution of ORNs to sleep. Interestingly, we observed little to no impact on sleep architecture when silencing *Ir8a*+ ORNs in virgin females, while the effects of silencing *Orco*+ were largely conserved (**Supplemental Figures 2-3**). Additionally, while many of the *Orco* and *Ir8a* nighttime effects observed in mated females (**Figure 1; Supplemental Figure 1**) were conserved in males, effects on daytime sleep were largely absent (**Supplemental Figures 4-5**). Finally, silencing *Ir25a*+ ORNs decreased nighttime sleep duration in males (**Supplemental Figure 4A, 4C**) and made them more hyperactive (**Supplemental Figure 5I-J**), a phenotype not previously observed in mated females (**Supplemental Figure 1**). Together, these results suggest that 1) ORN sleep modulation varies across internal states and in a cell type-specific manner, and 2) changes in sleep are likely attributed to changes in ORN excitability. Nonetheless, since silencing ORNs impacted sleep architecture most extensively in mated females (**Figure 1; Supplemental Figures 2-5**), we further investigated ORN sleep modulation within this context.

### Individual ORN types influence different parameters of day and nighttime sleep architecture

We next explored whether individual ORN types influence the same parameters of sleep architecture as their corresponding co-receptor populations, or if ensembles of ORNs mask the contributions of individual ORN types. Silencing *Orco*+ ORNs primarily consolidated nighttime sleep (**Figure 1**), whereas silencing *Ir8a*+ ORNs consolidated both day and nighttime. Together, these results suggested that the individual ORN types included in the *Ir8a-*GAL4 driver could modulate distinct temporal phases and parameters of sleep (**Figure 2Ai**). To test this hypothesis, we first combined the *Ir8a*-GAL4 with an *Orco*-GAL80^77^ to repress the expression of GAL4 in *Orco*+ ORNs and then reassessed the effects of BoNT-C on sleep (**Figure 2Aii**). This approach reduced the total number of glomeruli labeled by *Ir8a*-GAL4 from ∼11 to 9, specifically abolishing expression within the VC3 and VL2p glomeruli (**Figure 2B-C**). Silencing ORNs innervating the remaining glomeruli eliminated the impact on daytime sleep that was observed with full *Ir8a*+ expression, yet was still sufficient to increase total nighttime sleep duration (**Figure 2D-F**). Furthermore, nighttime sleep episode duration was increased while sleep episode number was decreased (**Figure 2J-L**), supporting that nighttime sleep architecture was still consolidated as observed when all *Ir8a*+ ORN types were silenced (**Figure 1**). Together, these results suggested that individual ORN types do indeed modulate specific temporal phases and parameters of sleep architecture.

**Figure 2.**
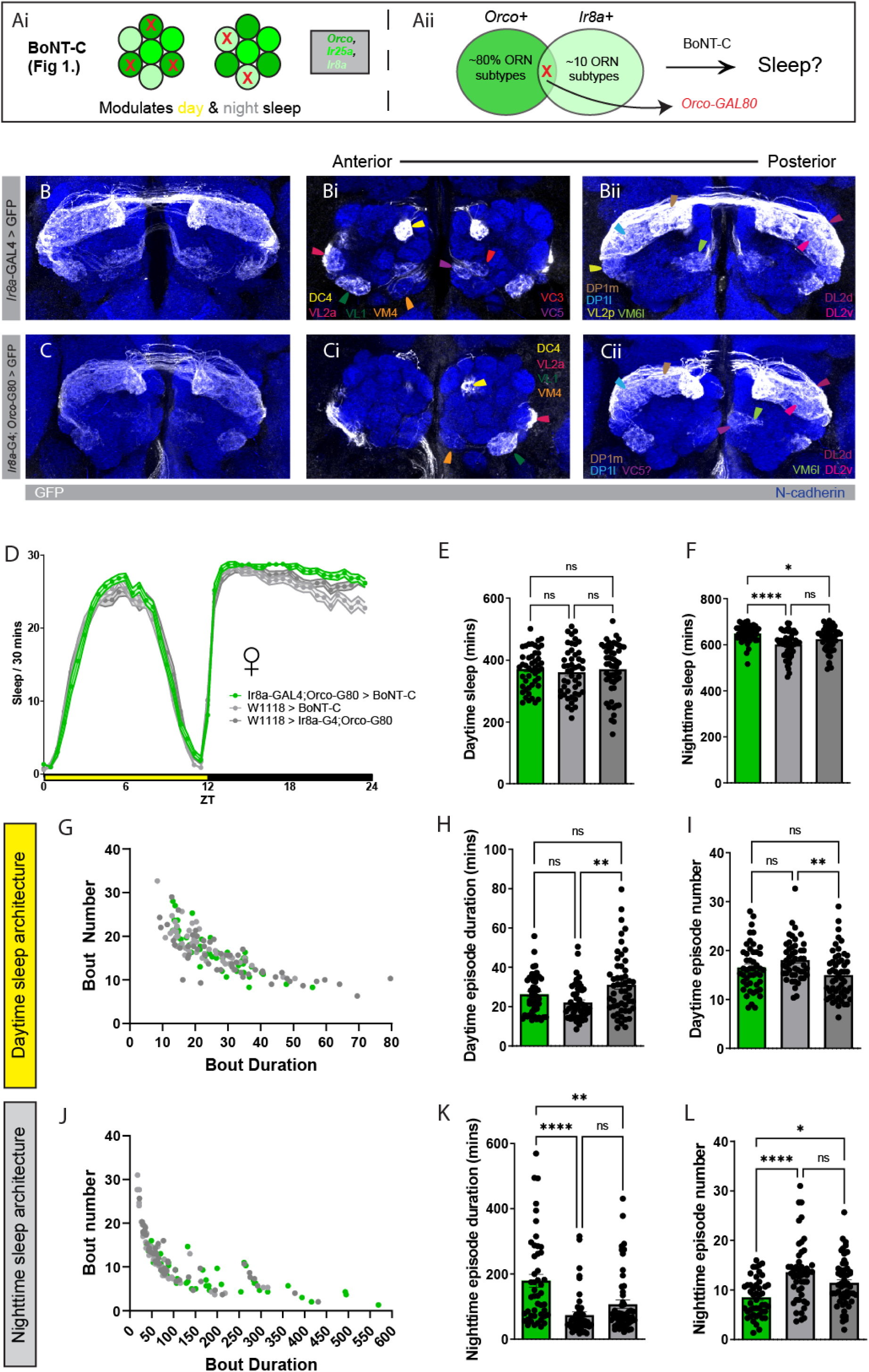
Altering the expression pattern of *Ir8a*-GAL4 reduces the effect of BoNT-C on daytime sleep. **A)** Schematic of hypothesis testing the contributions of individual ORN types on sleep. Since both the *Orco* and *Ir8a*-GAL4 drivers similarly influenced day and nighttime sleep architecture, we questioned whether silencing ORNs labeled in an altered expression pattern of *Ir8a*-GAL4 via an *Orco*-GAL80 would also alter the effects on sleep, relative to *Ir8a*-GAL4 > BoNT-C (**Figure 1**). **B-C**) Glomerular innervation patterns of ORNs labeled with *Ir8a*-GAL4 and *Ir8a*-GAL4;*Orco*-GAL80. Leftmost is whole-projection Z-stack images of the expression pattern of each driver line within the AL. Middle pains are anterior slices of the stack, while rightmost is posterior slices. GFP signal is driven in each driver line (white), while the neuropil are labeled by N-cadherin (blue). **D**) Sleep per 30 minutes of flies where ORNs labeled in an altered expression pattern of *Ir8a*-GAL4 are silenced via BoNT-C. Yellow bar represents daytime (ZT 0-12), while black is nighttime (ZT 12-24). **E-F**) Average sleep duration of **E**) daytime and **F**) nighttime sleep phases. **G**) Scatter plot of the per-fly daytime sleep architecture across each tested genotype. **H**) Average daytime sleep bout duration. **I**) Average daytime sleep bout number. **J**) Scatter plot of the per-fly nighttime sleep architecture. **K**) Average nighttime sleep bout duration. **L**) Average nighttime sleep bout number. In each behavior experiment, 3-7 day age mated female flies were tested, N=47-53 per genotype. Lightest gray = BoNT-C control, darker gray = Ir8a-GAL4;Orco-GAL80 control, green = *Ir8a*-GAL4;*Orco*-GAL80 > BoNT-C. See methods for details on statistical analysis. *p<0.05, **p<0.01, ***p<0.001, ****p<0.0001.

The *Ir8a*-GAL4;*Orco*-GAL80 experiments suggested that *Ir8a*+,*Orco*- ORNs are candidate nighttime sleep modulators, while ORNs innervating VC3 and VL2p glomeruli are candidate daytime sleep modulators (**Figure 3A**). To better understand the contributions of individual ORN types on sleep, we next performed a BoNT-C screen using drivers for the majority of the individual ORN types labeled within *Ir8a*-GAL4: *Ir31a*-GAL4 (innervates VL2p and DM3 glomeruli), *Ir41a*-GAL4 (VC5), *Ir64a*-GAL4 (DC4 and DP1m), *Ir75a*-GAL4 (DP1l), *Ir76a*-GAL4

**Figure 3.**
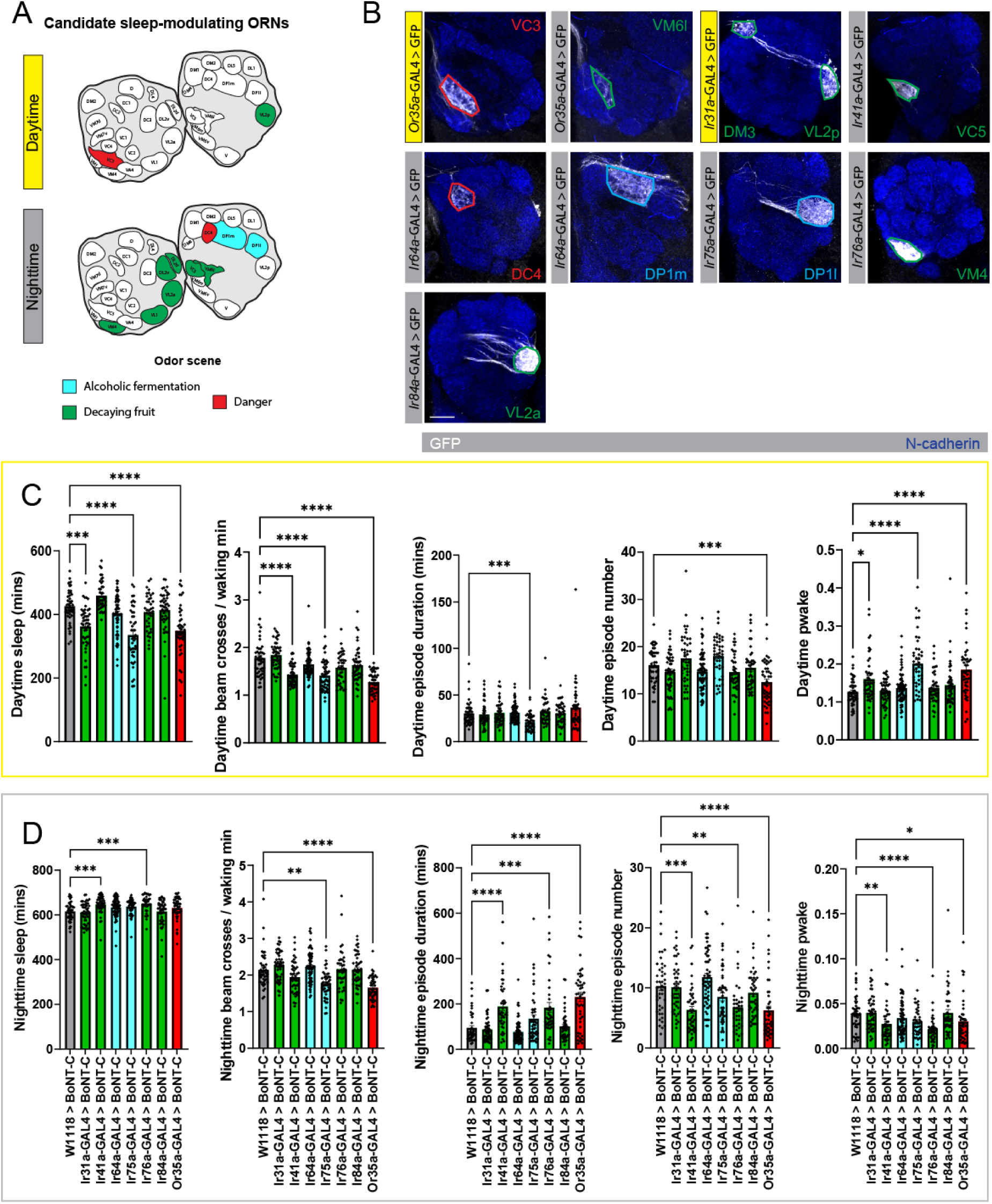
Neurotransmission of individual *Ir8a*+ ORN types shapes distinct parameters of day and nighttime sleep architecture. A) Schematic of candidate day and nighttime sleep-modulating *Ir8a*+ ORN types, inferred from experiments in **Figure 2**. Antennal lobe glomerular meshes adapted from Dr. Chris Potter’s lab website (https://potterlab.johnshopkins.edu/). Color of each glomeruli’s odor scene adapted from prior studies^42^ ^161^. **B**) Glomerular innervation patterns of individual *Ir8a*+ ORN type driver lines. Yellow nameplate marks candidate daytime sleep modulators, while gray marks nighttime. Glomerular boundaries are color-coded to match the odor scenes and the references used in **A**). GFP signal is driven in each GAL4 (white), while the neuropil is labeled by N-cadherin. Scale bar is 20µm. **C-D**) Daytime and nighttime sleep changes, respectively. Left to right: average sleep duration, average beam crosses per waking minute, average sleep bout duration, average sleep bout number, and average pwake. In each behavior experiment, 3-7 day age mated female flies were tested, N=39-63 per genotype. For **C-D**), the color of each experimental genotype matches the corresponding glomerular odor scene shown in **A-B**), while gray = BoNT-C control. See methods for details on statistical analysis. *p<0.05, **p<0.01, ***p<0.001, ****p<0.0001.

(VM4), *Ir84a*-GAL4 (VL2a), and *Or35a*-GAL4 (VC3 and VM6l) (**Figure 2B**; **Figure 3B**). Consistent with our hypothesis, only silencing three of the tested individual ORN types were sufficient to modulate daytime sleep (**Figure 3A**), although the effects did not directly phenocopy that of *Ir8a*-GAL4;*Orco*-GAL80 (**Figure 2**). Silencing *Ir31a*+ (VL2p, DM3), *Ir75a*+ (DP1l), and *Or35a*+ (VC3, VM6l) ORNs each decreased daytime sleep duration (**Figure 3C**), while silencing *Ir41a+* (VC5), *Ir75a+,* and *Or35a+* ORNs decreased activity per waking minute (**Figure 3C**). Only silencing *Ir75a*+ ORNs altered daytime sleep episode duration (**Figure 3C**), while silencing *Or35a+* ORNs were the only ORNs to alter daytime sleep episode number (**Figure 3C**). Finally, we found that silencing *Ir31a*+, *Ir75a*+, and *Or35a*+ ORNs also increased daytime pwake (**Figure 3C**). Overall, these results suggest that these individual ORNs modulate specific temporal phases and parameters of daytime sleep. However, since silencing individual *Ir8a*+ ORN types did not entirely match the daytime phenotypes observed for *Ir8a*-GAL4 (**Figure 1**), individual ORN types likely make distinct contributions to sleep rather than functioning in an additive manner.

In contrast to daytime, silencing individual *Ir8a*+ ORN types generally consolidated nighttime sleep architecture, similar to silencing *Ir8a*-GAL4 (**Figure 1**). Silencing *Ir41a*+ (VC5) and *Ir76a*+ (VM4) ORNs each increased total nighttime sleep duration (**Figure 3D**) via consolidated nighttime sleep architecture with increased sleep episode duration and decreased sleep episode number (**Figure 3D**). Interestingly, silencing *Or35a*+ (VC3 and VM6l) ORNs also consolidated nighttime sleep architecture independent of changes in total nighttime sleep duration (**Figure 3D**), however this and *Ir75a+* (DP1l) ORNs were the only types that affected nighttime locomotion (**Figure 3D**). Finally, silencing *Ir41a*+, *Ir76a*+, and *Or35a*+ ORN types each decreased nighttime pwake (**Figure 3D**), further supporting that silencing these individual ORN types consolidated nighttime sleep.

Overall, this screen of individual ORN types (**Figure 3**) reinforced properties of sleep modulation we observed in the initial co-receptor screen (**Figure 1)**: 1) Signaling from individual ORN types modulates specific temporal phases of sleep (day (*Ir31a* and *Ir75a*), nighttime (*Ir41a* and *Ir76a*), both (*Or35a*)). 2) Individual ORN types modulate different sleep parameters within the same temporal phase of sleep (silencing *Ir31a* increased the probability to awaken during the day, whereas silencing *Or35a* decreased the initiation of sleep during the day). However, screening individual ORN types also demonstrated that 3) Individual ORN types had distinct effects on sleep architecture relative to their co-receptor population (silencing *Ir8a+* ORNs consolidated both day and nighttime sleep architecture (**Figure 1**), but silencing individual ORN types that makeup *Ir8a*-GAL4 expression promoted daytime wake and nighttime sleep). Altogether, these results support a model in which interglomerular differences in ORN signaling shape distinct parameters of sleep architecture, as opposed to all ORNs having the same functions throughout the sleep/wake cycle.

### Serotonin receptor signaling across all ORN types differentially modulates day and nighttime sleep architecture

We next sought to identify candidate upstream signaling pathways of ORNs that could shape interglomerular differences of sleep modulation. Given the well-established role of serotonin (5-HT) in circadian rhythms and sleep in both vertebrates and invertebrates^78–82^, and broad expression of the serotonin 2B receptor (5-HT2B) by ORNs^83,84^, we hypothesized that this modulator could adjust the impact of each ORN type on sleep. Both scRNAseq datasets of Drosophila antennae^84^ and genomic tagging of 5-HT2B^85^ reveal that individual ORN types heterogeneously express 5-HT2B across glomeruli, suggesting that 5-HT signaling within the AL could shape stimulus-specific or ORN-specific effects on behavior, similar to how ORN signaling influences sleep (**Figures 1-3**).

To test the effects of 5-HT2B signaling in ORNs on sleep, we first used RNA interference (RNAi)^86,87^ to knockdown the expression of 5-HT2B in all ORNs via the *peb*-GAL4 driver^88^, which also expresses some auditory and gustatory sensory neurons (**Figure 4A**). To validate the effectiveness of the 5-HT2B RNAi, we leveraged flies with endogenously HA-tagged 5-HT2B^85^ and simultaneously expressed 5-HT2B RNAi in *peb*-GAL4 to assess changes in 5-HT2B expression. Knocking down 5-HT2B expression in *peb*-GAL4 removed nearly all 5-HT2B HA labeling within the AL and generally reduced the amount observed in the SEZ (**Figure 4B-C**), supporting that ORNs express the majority of 5-HT2B within the AL^83^ and confirming that this RNAi approach was sufficient to disrupt 5-HT2B expression. We then assessed the effects of knocking down 5-HT2B expression in *peb*-GAL4 on sleep. Knocking down 5-HT2B expression in *peb*-GAL4 decreased total daytime sleep duration and increased total nighttime sleep duration (**Figure 4D-F**), suggesting that 5-HT2B signaling across the neurons expressed within *peb*-GAL4 does not uniformly modulate sleep. Furthermore, daytime sleep duration was modulated independent of attributable changes in sleep architecture, which may have been due to behavioral variance of the control strains (data not shown). However, nighttime sleep was consolidated (**Figure 4G**) via increased nighttime sleep episode duration (**Figure 4H**), decreased number of sleep episodes (**Figure 4I**), and decreased nighttime pwake (**Figure 4J-L**), similar to the effects of silencing *Ir8a+* and *Orco*+ ORNs (**Figure 1**). Considering peb-GAL4 expresses other sensory neurons, it is difficult to directly attribute these observed changes in sleep to 5-HT2B signaling within ORNs. However, since auditory neurons do not seem to affect sleep ^30^ and gustatory neurons do in a diet-dependent manner ^89^, these results instead suggest that under baseline conditions reducing 5-HT2B expression within ORNs modulates day and nighttime sleep, consistent with silencing ORNs via BoNT-C (**Figures 1-3**) and the excitatory nature of 5-HT2B signaling^90–92^.

**Figure 4.**
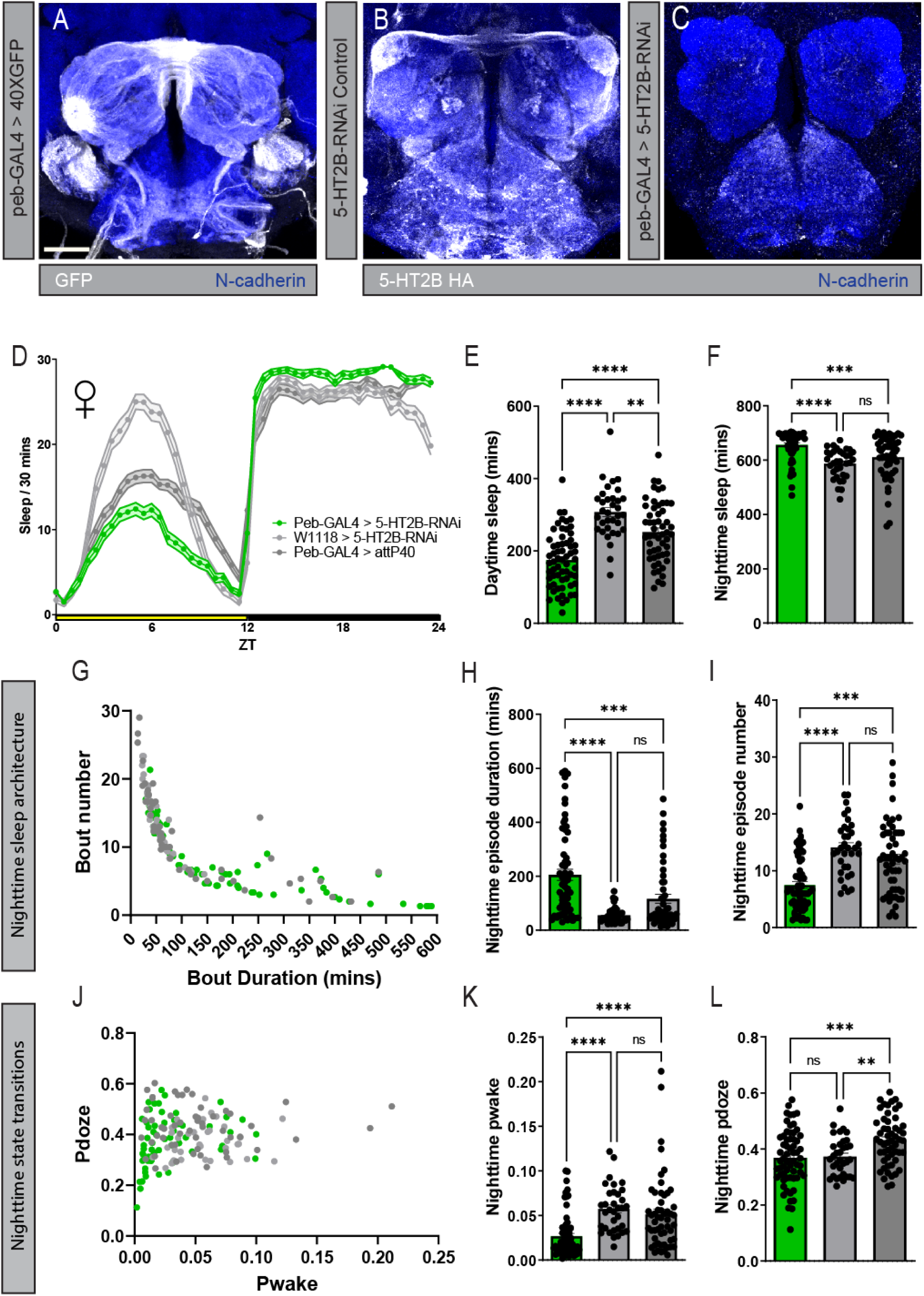
5-HT2B signaling across all ORN types differentially modulates day and nighttime sleep. **A)** Glomerular innervation pattern of *peb*-GAL4 driver line. GFP signal driven by the GAL4 (white), neuropil labeled by N-cadherin (blue). **B-C**) 5-HT2B HA labeling of the AL and SEZ in **B**) baseline conditions and **C**) animals with reduced 5-HT2B expression in all ORNs via *peb*-GAL4 > 5-HT2B-RNAi. 5-HT2B HA labeling (white) and neuropil labeled by N-cadherin (blue). Scale bar for **A-C**) is 30µm. **D**) Sleep per 30 minutes of flies where 5-HT2B expression is reduced in all ORNs via *peb*-GAL4 > 5-HT2B-RNAi. Yellow bar represents daytime (ZT 0-12), while black is nighttime (ZT 12-24). **E-F**) Average sleep duration of **E**) daytime and **F**) nighttime sleep phases. **G**) Scatter plot of the per-fly nighttime sleep architecture across each tested genotype. **H**) Average nighttime sleep bout duration. **I**) Average nighttime sleep bout number. **J**) Scatter plot of the per-fly nighttime state transition architecture across each genotype. **K**) Average nighttime pwake. **L**) Average nighttime pdoze. In each behavior experiment, 3-7 day age mated female flies were tested, N=34-62 per genotype. Lightest gray = 5-HT2B-RNAi control, darker gray = *peb*-GAL4 control, green = *peb*-GAL4 > 5-HT2B-RNAi. See methods for details on statistical analysis. *p<0.05, **p<0.01, ***p<0.001, ****p<0.0001.

### 5-HT2B signaling in distinct ORN co-receptor populations influences different parameters of day and nighttime sleep architecture

Our next question was whether 5-HT2B signaling in distinct ORN types modulated day and nighttime sleep architecture. The extent to which 5-HT2B signaling uniformly modulates ORNs is unclear as the expression of 5-HT2B in ORNs has only been quantified in a subset of ORNs^84,85^. We therefore used the publicly available SCope scRNA-seq antennal dataset^93,94^ to quantify the expression of 5-HT2B within each individual ORN type and organized them into their corresponding co-receptor populations. Consistent with previous studies^83^, this approach revealed that the 5-HT2B receptor is expressed in nearly every ORN class (**Figure 5A**). Furthermore, many individual ORN types with the highest expression levels were consistent across studies, such as *Or67d*+ DA1-innervating ORNs^84,85^, suggesting this approach was sufficiently sensitive to compare relative levels of 5-HT2B expression levels between ORN types. In general, *Orco+* ORNs had more 5-HT2B receptor expression than ORNs from other co-receptor classes, but individual ORN types from *Ir8a* and *Ir25a* co-receptor populations also displayed similar high levels of expression (**Figure 5A**). Additionally, there was no relationship between the level of 5-HT2B expression and the conveyed odor scene of ORNs. Taken together, these results suggest that distinct ORN types from a variety of co-receptor families may be modulated by 5-HT within the context of sleep **(Figure 5B**).

**Figure 5.**
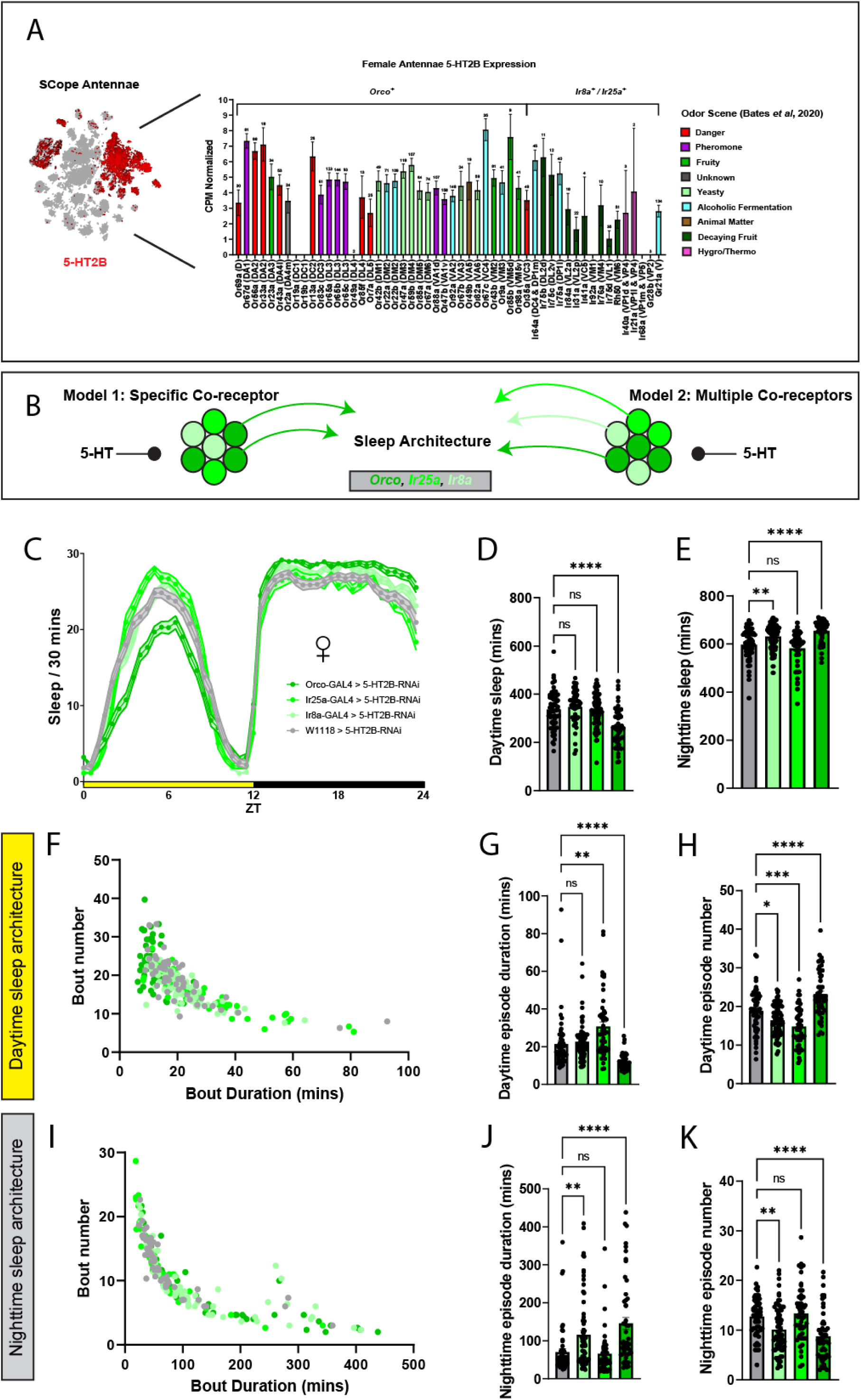
5-HT2B signaling across ORN co-receptor populations differentially modulates day and nighttime sleep. **A)** SCope female antennae scRNA-seq readouts, average 5-HT2B expression across individual ORN types. Odor scenes of individual ORN types and color schemes adapted from ^42^ and ^161^. **B**) Schematic of opposing models where either a distinct ORN co-receptor population, such as *Orco*+ ORNs, are the only types modulated by 5-HT within the context of sleep vs one in which the 5-HT modulation of ORNs from various co-receptor populations influence sleep. **C**) Sleep per 30 minutes of flies where 5-HT2B expression is reduced in different ORN co-receptor populations via 5-HT2B-RNAi. Yellow bar represents daytime (ZT 0-12), while black is nighttime (ZT 12-24). **D-E**) Average sleep duration of **D**) daytime and **E**) nighttime sleep phases. **F**) Scatter plot of the per-fly nighttime sleep architecture across each tested genotype. **G**) Average nighttime sleep bout duration. **H**) Average nighttime sleep bout number. **I**) Scatter plot of the per-fly nighttime state transition architecture across each genotype. **J**) Average nighttime pwake. **K**) Average nighttime pdoze. In each behavior experiment, 3-7 day age mated female flies were tested, N=51-70 per genotype. Gray = 5-HT2B-RNAi control, lightest green = *Ir8a*-GAL4 > 5-HT2B-RNAi, brighter green = *Ir25a*-GAL4 > 5-HT2B-RNAi, darkest green = *Orco*-GAL4 > 5-HT2B-RNAi. See methods for details on statistical analysis. *p<0.05, **p<0.01, ***p<0.001, ****p<0.0001.

Indeed, knockdown of 5-HT2B in each ORN co-receptor population modulated different parameters of day and nighttime sleep (**Figure 5C**), indicating that 5-HT2B signaling in ORNs from more than one co-receptor population can impact sleep. For instance, knockdown of 5-HT2B expression within *Orco*+ ORNs decreased total daytime sleep duration and increased total nighttime sleep duration (**Figure 5D**), most similar to knockdowns within *peb*-GAL4 (**Figure 4**), whereas knockdown of 5-HT2B in *Ir8a*+ ORNs only increased nighttime sleep duration and knockdown in *Ir25a*+ ORNs had no effect on sleep duration (**Figure 5D-E**). Furthermore, knockdown of 5-HT2B expression in each co-receptor population differentially modulated day and nighttime sleep architecture (**Figure 5F-K**). Knocking down 5-HT2B expression within *Orco*+ ORNs fractured daytime sleep (**Figure 5F**) by decreasing daytime sleep episode duration (**Figure 5G**) and increasing episode number (**Figure 5H**), while nighttime sleep was instead consolidated (**Figure 5I-K**). 5-HT2B knockdown in *Ir25a*+ ORNs consolidated daytime sleep architecture independent of affecting daytime sleep duration (**Figure 5F-H**), while knockdown within *Ir8a*+ ORNs consolidated nighttime sleep, similar to *Orco*+ ORNs (**Figure 5I-K**).

These results support a model in which heterogeneous 5-HT2B expression across ORN types differentially shapes day and nighttime sleep architecture. This is also consistent with our findings that different co-receptor populations modulate distinct temporal phases of sleep (**Figure 1**). Furthermore, 5-HT2B signaling within different co-receptor populations again modulates distinct parameters of sleep within a given temporal phase, rather than having a uniform effect on day and nighttime. For example, knocking down 5-HT2B expression within *Orco+* ORNs fractured daytime sleep but consolidated nighttime, while knockdowns in *Ir8a+* ORNs only consolidated nighttime sleep. Finally, since 5-HT2B knockdowns in some ORN co-receptor populations do not entirely phenocopy their corresponding BoNT-C phenotypes (**Figure 1**), it is indicative that 1) the role of each ORN type in sleep is not caused, but rather modulated by 5-HT, and 2) ORNs may receive more than one modulatory input within sleep contexts. Overall, these findings reinforce that different ORN types modulate distinct parameters of sleep architecture and 5-HT2B signaling in part shapes these responses.

### 5-HT2B signaling in individual ORN types heterogeneously modulates sleep architecture

We next considered whether 5-HT2B signaling in individual ORN types promotes differences in sleep modulation relative to their co-receptor populations (**Figure 5**), as was observed in the BoNT-C experiments (**Figures 2-3**). The reduction of 5-HT2B expression within *Orco* and *Ir8a*-GAL4 both consolidated nighttime sleep architecture (**Figure 5**). 5-HT2B knockdowns within *Ir8a*-GAL4;*Orco*-GAL80 flies also increased nighttime sleep duration and consolidated nighttime sleep architecture (**Supplemental Figure 6**), suggesting that inhibiting 5-HT2B signaling within both *Ir8a*+ or *Ir8a*-GAL4;*Orco*-GAL80+ ORN types (**Figure 2C**) had a net consolidating effect in the nighttime (**Figure 6A**). Individual *Ir8a*+ ORN types each expressed 5-HT2B, as validated by simultaneously labeling glomerular boundaries with GFP and HA-tagged 5-HT2B (**Figure 6B**). Interestingly, 5-HT2B knockdowns in individual *Ir8a*+ ORN types modulated both day and nighttime sleep architecture, as opposed to having a uniform effect on nighttime sleep (**Figure 6C**). For instance, *Ir75a*+ (DP1l) ORNs were the only ORN type in which a 5-HT2B knockdown modulated daytime sleep, causing a decrease in total sleep duration (**Figure 6C**). Knockdowns in *Ir41a*+ (VC5), and *Or35a*+ (VC3, VM6l) ORNs decreased daytime activity per waking minute, while *Ir64a*+ (DC4, DP1m) ORNs increased it (**Figure 6C**), suggesting some changes in daytime sleep may be attributed to changes in locomotion. However, parameters of daytime sleep architecture were also modulated by 5-HT2B knockdowns in multiple ORN types. 5-HT2B knockdowns in *Ir31a+* (VL2p and DM3)*, Ir41a+,* and *Ir75a+* ORNs decreased daytime sleep episode duration (**Figure 6C**), while knockdowns in *Ir64a*+ and *Ir84a*+ (VL2a) ORNs instead decreased daytime sleep episode number (**Figure 6C**). Finally, knockdown in *Ir41a*+, *Ir75a*+, and *Or35a*+ ORNs each increased daytime pwake (**Figure 6C**), altogether suggesting that reducing 5-HT2B signaling in individual *Ir8a*+ ORN types generally fractures sleep to promote wakefulness in the daytime, in contrast to the effects observed from knockdowns in *Ir8a*-GAL4 (**Figure 5**).

**Figure 6.**
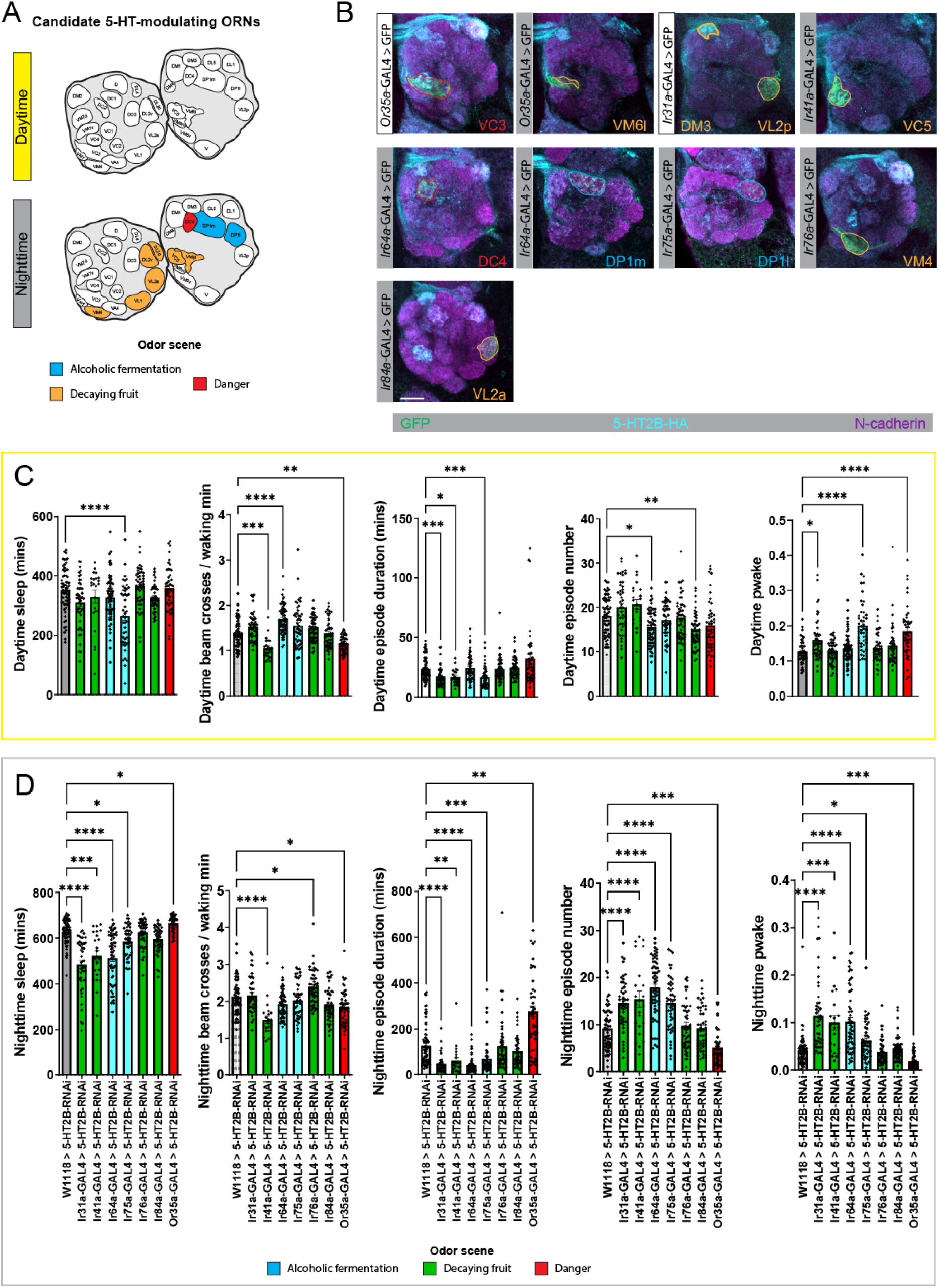
5-HT2B signaling within individual *Ir8a*+ ORN types modulates distinct parameters of day and nighttime sleep architecture. A) Schematic of candidate *Ir8a*+ ORN types modulated by 5-HT within sleep contexts, inferred from experiments in **Supplemental Figure 6**. Antennal lobe glomerular meshes adapted from Dr. Chris Potter’s lab website (https://potterlab.johnshopkins.edu/). Color of each glomeruli’s odor scene adapted from prior studies^42^ ^161^. **B**) Glomerular innervation patterns of individual *Ir8a*+ ORN type driver lines while co-labeling for endogenous HA-tagged 5-HT2B. White nameplate marks ORN types with no hypothesized effect from 5-HT2B knockdowns, while gray marks candidate nighttime modulators. Glomerular boundaries are color-coded to match the odor scenes and the references used in **A**). GFP signal is driven in each GAL4 (green), HA-tagged 5-HT2B (cyan), and the neuropil is labeled by N-cadherin (magenta). Scale bar is 20µm. **C-D**) Daytime and nighttime sleep changes, respectively. Left to right: average sleep duration, average beam crosses per waking minute, average sleep bout duration, average sleep bout number, and average pwake. In each behavior experiment, 3-7 day age mated female flies were tested, N=22-63 per genotype. For **C-D**), the color of each experimental genotype matches the corresponding glomerular odor scene shown in **D**), while gray = 5-HT2B-RNAi control. See methods for details on statistical analysis. *p<0.05, **p<0.01, ***p<0.001, ****p<0.0001.

5-HT2B knockdowns in individual *Ir8a*+ ORN types heterogeneously modulated nighttime sleep duration. 5-HT2B knockdowns in *Ir31a+* (VL2p, DM3)*, Ir41a+* (VC5), *Ir64a+* (DC4, DP1m), and *Ir75a+* (DP1l) ORNs decreased total nighttime sleep duration while knockdown in *Or35a*+ (VC3, VM6l) ORNs increased it (**Figure 6D**). 5-HT2B knockdown in *Ir41a*+ and *Or35a*+ ORNs decreased nighttime activity per waking minute, while *Ir76a*+ (VM4) ORNs weakly increased it (**Figure 6D**), suggesting the majority of nighttime phenotypes were largely independent of changes in locomotion. Nighttime sleep architecture was also modulated by 5-HT2B knockdowns in multiple ORN types. Knockdown in *Ir31a*+, *Ir41a*+, *Ir64a*+, and *Ir75a*+ ORNs each fractured nighttime sleep via decreased nighttime sleep episode duration (**Figure 6D**) and increased sleep episode number (**Figure 6D**). In contrast, knockdown in *Or35a*+ ORNs consolidated nighttime sleep architecture via increased nighttime sleep episode duration (**Figure 6D**) and decreased nighttime sleep episode number (**Figure 6D**). Finally, 5-HT2B knockdowns in *Ir31a*+, *Ir41a*+, *Ir64a*+, and *Ir75a*+ ORNs each increased nighttime pwake, while knockdown in *Or35a*+ ORNs decreased it (**Figure 6D**), altogether demonstrating that 5-HT2B signaling within individual *Ir8a*+ ORN types maintains different parameters of nighttime sleep architecture.

This screen revealed that the effects of 5-HT2B signaling on sleep within this network are complex, with ORN types differing in terms of the sleep phase that they impact. The effects of 5-HT2B signaling in *Ir8a*+ ORNs were also generally stronger at nighttime, consistent with our previous findings (**Figure 5; Supplemental Figure 6**). However, since 5-HT2B knockdowns in all ORNs and *Orco*+ ORNs reduced daytime sleep (**Figure 4-5**), there are likely other unidentified ORNs more strongly modulated in the daytime. Importantly, these results support a model in which 5-HT2B signaling of distinct ORN types heterogeneously modulates day and nighttime sleep architecture, similar to the 5-HT2B knockdowns in different ORN co-receptor populations (**Figure 5**) and silencing ORNs via BoNT-C (**Figures 1-3**).

### CSDn 5-HT signaling predominantly modulates nighttime sleep architecture

Our final question was whether synaptic 5-HT release within the AL modulated sleep architecture in a similar fashion as 5-HT2B knockdowns in ORNs (**Figures 4-6**). The AL receives synaptic 5-HT from a single pair of cells known as the contralaterally-projecting serotonin-immunoreactive deutocerebral neurons (the “CSDns”)^95,96^. The CSDns provide input to nearly every olfactory cell class within the AL and to higher-order networks within the olfactory pathway^97^. Importantly, the CSDns differentially innervate the glomeruli of the AL^95^ and share non-uniform connectivity with each ORN type^97^, suggesting that CSDn 5-HT signaling could heterogeneously modulate ORN responses. Additionally, the CSDns are an evolutionarily conserved serotonin cell class across many invertebrate species^96,98,99^ and 5-HT concentrations in the ALs of other invertebrates that possess CSDns fluctuate throughout the day^100^. Altogether, these findings suggest that CSDn 5-HT signaling may have time-dependent effects on behavior and are one candidate modulator of ORNs within the context of sleep.

To characterize the contributions of the CSDns on sleep, we first tested the effects of CSDn 5-HT release on sleep. Vesicular monoamine transporter (*vmat*) packages 5-HT and other monoamines into vesicles for synaptic release^101^. Therefore, we hypothesized that reducing its expression within the CSDns, and subsequently reducing CSDn 5-HT release, would disrupt sleep. We first addressed this question by driving the expression of RNAi targeting *vmat* and Dicer-2 within a broad CSDn driver (R60F02-GAL4)^102–104^ (**Supplemental Figure 7J**), which eliminates all 5-HT immunolabeling within the AL^85^ (**Figure 7A-B**). Constitutively inhibiting CSDn 5-HT release predominantly modulated nighttime sleep architecture independent of daytime (**Figure 7C-K**), consistent with many of the effects of 5-HT2B knockdowns in ORNs (**Figures 4-6**). Total nighttime sleep duration was increased relative to controls (**Figure 7E**) and nighttime sleep architecture was consolidated (**Figure 7I**) via increased sleep episode duration (**Figure 7J** and decreased sleep episode number (**Figure 7K**). As a secondary form of validation, we repeated these experiments using a stringent CSDn driver (MB465C-spGAL4)^95,105^ (**Supplemental Figure 7K**) and found similar trending effects on sleep, although at a reduced magnitude (**Supplemental Figure 7A-I**). Upon closer inspection of both driver line expression patterns, we found that MB465C-spGAL4 stochastically labeled CSDns within the brain compared to R60F02-GAL4 (**Supplemental Figure 7J-L**). We therefore attributed these differences in behavior to the strength of *vmat*-RNAi affecting both CSDns and continued using R60F02-GAL4 for further experimentation.

**Figure 7.**
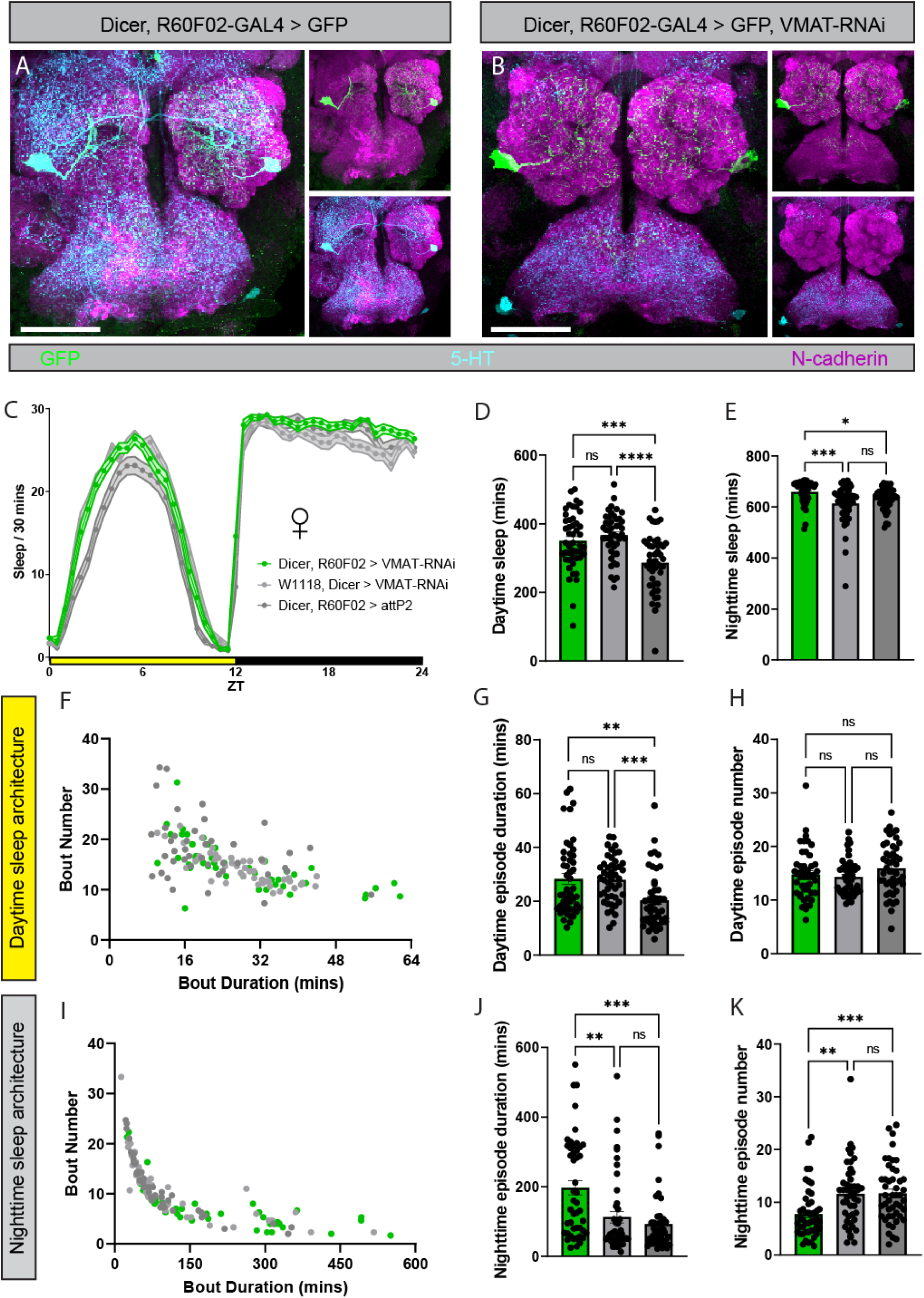
CSDn 5-HT release predominantly modulates nighttime sleep architecture. A-B) Glomerular innervation patterns of CSDns under **A**) baseline conditions and **B**) animals where CSDn 5-HT signaling is inhibited via Dicer, R60F02-GAL4 > VMAT-RNAi. GFP signal is driven in the R60F02-GAL4 (green), 5-HT antibody (cyan), and the neuropil is labeled by N-cadherin (magenta). Scale bar is 30µm. **C**) Sleep per 30 minutes of flies where CSDn 5-HT signaling is inhibited via Dicer, R60F02-GAL4 > VMAT-RNAi. Yellow bar represents daytime (ZT 0-12), while black is nighttime (ZT 12-24). **D-E**) Average sleep duration of **D**) daytime and **E**) nighttime sleep phases. **F**) Scatter plot of the per-fly daytime sleep architecture across each tested genotype. **G**) Average daytime sleep bout duration. **H**) Average daytime sleep bout number. **I**) Scatter plot of the per-fly nighttime sleep architecture across each genotype. **J**) Average nighttime sleep bout duration. **K**) Average nighttime sleep bout number. In each behavior experiment, 3-7 day age mated male flies were tested, N=47 per genotype. Lightest gray = Dicer and VMAT-RNAi control, darker gray = Dicer, R60F02-GAL4 control, green = Dicer, R60F02-GAL4 > VMAT-RNAi. See methods for details on statistical analysis. *p<0.05, **p<0.01, ***p<0.001, ****p<0.0001.

Since co-transmission from serotonergic neurons modulates sleep in *Drosophila*^106^, and there are a variety of reports suggesting the CSDns co-transmit, we next sought to characterize the effects of different signaling mechanisms utilized by the CSDns on sleep. First, a recent study suggests that the CSDns release dopamine^107^, therefore *vmat* knockdowns in the CSDns (**Figure 7**) may also affect CSDn dopamine signaling. As a secondary form of validation that CSDn 5-HT modulates sleep, we tested whether CSDn 5-HT transporter (*SerT)* signaling modulates sleep architecture. *SerT* functions within serotonergic neurons to re-uptake 5-HT from the synapse, limiting the duration of the effects of 5-HT on its downstream targets^108^. Importantly, genome-wide knockout of *SerT* increases total daytime and nighttime sleep duration^109^, suggesting *SerT* functionally impacts sleep in flies, but the contributions of the CSDns on these responses are unknown. To address this gap in knowledge, we disrupted CSDn *SerT* function by driving the expression of a *SerT* dominant negative (*SerT*-DN)^110^ within the R60F02-GAL4. Blocking 5-HT reuptake via *SerT*-DN reduces 5-HT levels of the presynaptic neuron^110^, therefore we hypothesized that inhibiting CSDn *SerT* signaling would similarly consolidate nighttime sleep as in *vmat*-RNAi (**Figure 7**). Indeed, constitutively inhibiting CSDn *SerT* signaling increased nighttime sleep duration and consolidated nighttime sleep architecture (**Supplemental Figure 8**), in addition to weakly consolidating daytime sleep. Altogether, these results further supported that CSDn 5-HT signaling modulates nighttime sleep architecture.

Next, although there are conflicting reports that the CSDns co-transmit acetylcholine (ACh)^104,111^, we questioned whether CSDn ACh signaling affected sleep. To test this hypothesis, we drove the expression of a previously validated RNAi targeting choline acetyltransferase (*ChAT*)^112^ and Dicer-2 within the R60F02-GAL4, however we did not observe any consistent changes in sleep duration or sleep architecture relative to controls (**Supplemental Figure 9**). Finally, we tested whether the CSDns were peptidergic and if this contributed to sleep. We first leveraged a combination of GAL4 drivers and mRNA labeling via hybridization chain reaction^113–115^ to determine if the CSDns expressed genes related to peptide signaling, such as *dimm*^116^, *syt-ɑ*^117^, and *stacl*^118,119^ (**Supplemental Figure 10A-C**). We found weak expression of *stacl* puncta within the boundaries of the CSDn cell bodies (**Supplemental Figure 10B**), suggesting that the CSDns may be *stacl+* and therefore could be peptidergic. We therefore tested whether decreasing the expression of *stacl* in the CSDns modulated sleep by driving the expression of *stacl*-RNAi in the R60F02-GAL4. Through this approach, we did not detect any consistent changes in sleep architecture relative to controls (**Supplemental Figure 10D-L**). Overall, the results of each of these experiments suggest that CSDn 5-HT release within olfactory circuits is important during the context of nighttime sleep, while other sources of 5-HT likely modulate them in the daytime.

## Discussion

In this study, we provide new insights into the mechanisms by which the olfactory system influences sleep architecture in *Drosophila melanogaster*. Consistent with previous studies^8,30^, our findings position olfactory receptor neurons (ORNs) as a key cell type within olfactory networks that shape sleep architecture. However, we report that the neurotransmission of distinct ORN types influence different temporal phases of sleep and parameters of sleep architecture, rather than all ORNs having a uniform influence on sleep. Furthermore, we find that broad metabolic needs and specific neuromodulatory pathways can alter the influence of distinct ORN types on sleep, altogether supporting a model in which the influence of distinct odor channels can be up and down-regulated to adjust the depth of sleep.

### ORN-specific modulation of sleep architecture

The overall contribution of ORNs to sleep regulation are complex and by no means uniform. Multiple studies suggest that removal of the third antennal segment increases total sleep duration, but these responses seem to be attributed to post-injury inductions of sleep via glial signaling^28,29,120^. A separate study showed that this same ablation instead decreases total sleep duration, but this may have been due to age-related differences in injury induction or diet fed to flies^30^. Interestingly, the same study reports that thermogenetically silencing *Orco*+ ORNs or gluing the third antennal segment to block olfactory transduction also decreases sleep^30^, suggesting that broadly inhibiting ORN signaling promotes wakefulness separate from injury. Additionally, genetic knockouts of genes related to olfactory transduction such as *Orco* differentially modulates day and nighttime sleep in male and female flies^32^. Altogether, these studies support that the influence of ORNs seemingly depends on the mechanism by which they are dysregulated and the internal context of the animal.

Here, we find that constitutively silencing distinct ORN types (and thereby distinct olfactory pathways) modulates different temporal phases and parameters of sleep architecture (**Figure 1**). While no other study to our knowledge has investigated the contributions of *Ir8a+* or *Ir25a*+ ORNs on sleep, our results are similar to *Orco* genetic knockouts^32^ in that constitutively silencing *Orco*+ ORNs also differentially modulates day and nighttime sleep. However, knockouts of ORN co-receptors are known to reduce the overall number of cells within distinct ORN types^121^, thus some differences between sleep phenotypes of *Orco*-GAL4 > BoNT-C (this study) and *Orco* knockout flies may be due to cell death. Additionally, silencing ORNs via BoNT-C alters sleep differently from that of the net decrease in total sleep duration observed from antennectomy or conditional silencing approaches^30^. Considering the age range, diet, sex, and isogenizing methodologies used for the flies were similar across both studies, conditionally removing olfactory signals may produce more uniform effects on sleep compared to constitutive silencing. However, since temperature is processed in the AL^63^, it is also possible that thermogenetically manipulating ORNs produces a confounding effect on sleep. In this study, we report that silencing the *Ir25a*-GAL4 (which includes temperature-sensing cells and other sensory cells across the body^61,65^) had little effect on baseline sleep architecture. However, considering *Ir25a*+ neurons are required for temperature synchronization of the circadian clock^122^, some ORN sleep phenotypes may change in a temperature-dependent manner. To avoid these concerns, future research utilizing conditional approaches to investigate the effects of ORN dysfunction on sleep could instead leverage optogenetic tools^123^.

We also report that silencing large populations of ORNs masks the contributions of individual ORN types, as was the case with *Ir8a*+ ORNs (**Figures 2-3**). This may indicate a combinatorial model of sleep modulation, where one individual ORN type may increase sleep and another decrease it, thus altering both simultaneously could have no net effect on sleep. However, a combinatorial model may also suggest that altering two ORN types with the same function in sleep would enhance the overall effect. Instead, we observed that the magnitude at which olfactory impairment influences sleep does not always scale to the number of different ORN types being dysregulated. For example, silencing the majority of ORNs via *Orco*-GAL4 (**Figure 1**) consolidates nighttime sleep just as strongly as silencing some individual ORN types (**Figure 3**). Additionally, silencing ORNs conveying a variety of odor scenes modulate day and nighttime sleep in similar fashions, as opposed to, for example, only danger or food-related ORNs being relevant within the context of sleep (**Figure 3**). Taken together, these findings suggest that changes in distinct ORN or olfactory pathways modulate sleep architecture, potentially representing a form of degeneracy within the olfactory circuit, especially in nighttime sleep contexts. Under baseline conditions, a degenerate model of olfactory modulation of sleep ensures the animal can appropriately awaken in the presence of a variety of odor contexts as opposed to the network being reliant on the cues of only a few channels. Alternatively, when these pathways are constitutively inhibited in different patterns, this heterogeneously modulates sleep architecture, suggesting that downstream connections within these pathways flexibly shape sleep architecture. Therefore, it is likely that the net weight or value of these signals within sleep contexts may be represented at the level of higher order neurons within the olfactory circuit not tested in this study, such as wake-promoting dopaminergic PPL1/PAM cells whose baseline activity levels reflect the sleep/wake state of the fly^9^.

### Neuromodulation of olfactory signaling within the context of sleep

In this study, we also determined that reducing the expression of the 5-HT2B receptor in ORNs heterogeneously modulates baseline sleep architecture (**Figures 4-6**), reflective of many phenotypes observed using BoNT-C (**Figures 1-3**) and consistent with a degenerate model of olfactory modulation of sleep. This is consistent with previous work broadly linking 5-HT^81,124,125^ and 5-HT2B signaling^126^ to sleep in *Drosophila*, but suggests a new function of 5-HT2B within the olfactory circuit. However, since ORNs are known to express a variety of aminergic and peptidergic receptors^73–76,84,127–130^, 5-HT is likely just one of many modulators of ORNs within the context of sleep.

One mechanism in which degeneracy can be promoted at a network level is if the sub-circuits involved are only activated in specific contexts^131–133^. The 5-HT2B receptor acts upon the Ca^2+^ and IP3 pathway to increase neuronal excitability^90–92^, therefore it is likely that endogenous 5-HT enhances the influence of ORNs on sleep. Interestingly, our results suggest that 5-HT does not uniformly affect ORNs due to the heterogeneous expression levels of 5-HT2B across ORN types. Consistent with previous studies^84,85^, we show that the expression of 5-HT2B mRNA transcripts varies across individual ORN types (**Figure 5A**) suggesting that individual ORN pathways may be more susceptible to 5-HT signaling than others. It may even be possible that the expression of 5-HT2B in given ORN types changes depending on the internal and external demands of the fly, similar to what has been shown in other receptor pathways expressed by ORNs^67,74,75^. This is in line with our results suggesting the influence of *Ir8a*+ ORN types shifts across the mating status of the fly (**Supplemental Figures 2-5**). However, since olfactory PNs and local interneurons (LNs) also express 5-HTRs^83^, it is possible interglomerular differences in sleep regulation are promoted by a combination of direct and indirect 5-HT modulation of ORN-PN connections.

Here, we also establish the CSDns as a sleep-modulating 5-HT cell type and one candidate source of 5-HT to ORNs within the context of sleep (**Figure 7**). Inhibiting CSDn 5-HT release predominantly consolidates nighttime sleep architecture, consistent with many phenotypes observed via 5-HT2B knockdowns in ORNs (**Figures 4-6**). However, since it is thought that ORNs and other olfactory cells receive paracrine 5-HT signaling from the hemolymph ^104,134^, there are likely unidentified cellular sources of 5-HT to ORNs. Our results support this notion, as inhibition of 5-HT release by the CSDns did not reproduce the effects of 5-HT2B knockdown in ORNs on daytime sleep (**Figure 4, 5, 7**) . Nonetheless, these results suggest that the CSDns are likely to be wake-promoting under baseline conditions. Since broader manipulations of the serotonin system promote sleep^135,136^, these results suggest distinct 5-HT networks instead heterogeneously modulate sleep/wake, consistent with prior findings in *Drosophila*^106,126,136,137^ and vertebrates^138–142^.

Interestingly, serotonergic “dorsal pair medial” (DPM) neurons, which innervate the mushroom bodies, specifically promote nighttime sleep^106,109,143^. Taken together with the findings from this study, synaptic 5-HT signaling within primary and higher order stages of olfactory processing convergently modulates nighttime sleep, however it is unlikely that CSDns and DPMs modulate the same circuits within the context of sleep. DPMs target neurons innervating the mushroom body lobes^144^, while the CSDns are the sole source of synaptic 5-HT within the AL, mushroom body calyx, and lateral horn^95^. Additionally, the CSDns and DPMs have different functions within olfaction, in that DPMs are required for olfactory memory consolidation^106,143,145–147^, while CSDns primarily impact olfactory sensitivity and detection^103,104,148–150^. Thus, it is possible that DPM 5-HT release consolidates nighttime sleep in an experience-dependent manner, while CSDn 5-HT release instead fractures nighttime sleep and may even adjust the probability of awakening in the presence of odors of interest.

Altogether, this work supports a model in which ORNs act collectively to shape sleep architecture, but their recruitment can be up and downregulated by the physiological state of the animal. Thus, individual ORN types are subsequently differentially modulated and the apparent degeneracy within the system could be a means by which different contexts impact sleep.

## Acknowledgements

This work was funded by NIH DC-016293 (AMD), NSF IOS 2114775 (AMD) and NIH T32 GM132494 (OMC). We thank the Bloomington *Drosophila* Stock Center for their invaluable service which is supported by NIH P40 OD018537. We would also like to thank Dr. Chris Vecsey for assistance using SCAMP, Dr. Annika Barber and Dr. Divya Sitaraman for experimental feedback, in addition to Hazem Attal, Abigail Long, Maggie Robertson, Farzaan Salman, Marryn Bennett, Ethan Mick, and Tessa Cessario for technical support. Finally, we thank Dr. Matthew Kayser for his feedback on earlier versions of this manuscript and Dr. Dion Dickman, Dr. Quentin Gaudry, Dr. David Krantz, and Dr. Henrike Scholz for providing fly lines.

## Methods

### Fly stocks

All fly stocks were raised on standard cornmeal/agar/yeast medium at 24°C on a 12:12 light/dark cycle at 50-60% humidity. Flies of 3-7 days of age were used for every experiment. To control for effects of genetic background and presence of balancer chromosomes, the majority of the GAL4 and UAS lines used in behavioral experiments were backcrossed 5 generations with iso31 w[1118] flies^151^ or two generations with a iso31 w[1118]; Cyo/Sco;Mkrs/TM6B stock and then selected against balancers for a minimum of two more generations prior to use. RNAi experiments tested females of controlled heterozygous genetic backgrounds and males with iso31 backgrounds relative to RNAi or RNAi insertion site controls. When expressing Dicer-2 to increase the RNAi efficacy^152,153^, stable lines were generated by crossing GAL4s of interest to a iso31 w[1118],UAS-Dicer2;Cyo/Sco;Mkrs/TM6B stock for two generations and then similarly selecting against balancers as before. In behavior experiments screening multiple GAL4 lines at once, we chose to compare experimental groups to the w[1118] background control expressing the UAS tool of interest, similar to other studies^154–158^. In behavior experiments testing one GAL4, we compared experimental groups to respective GAL4 and UAS controls.

**Table 1.**
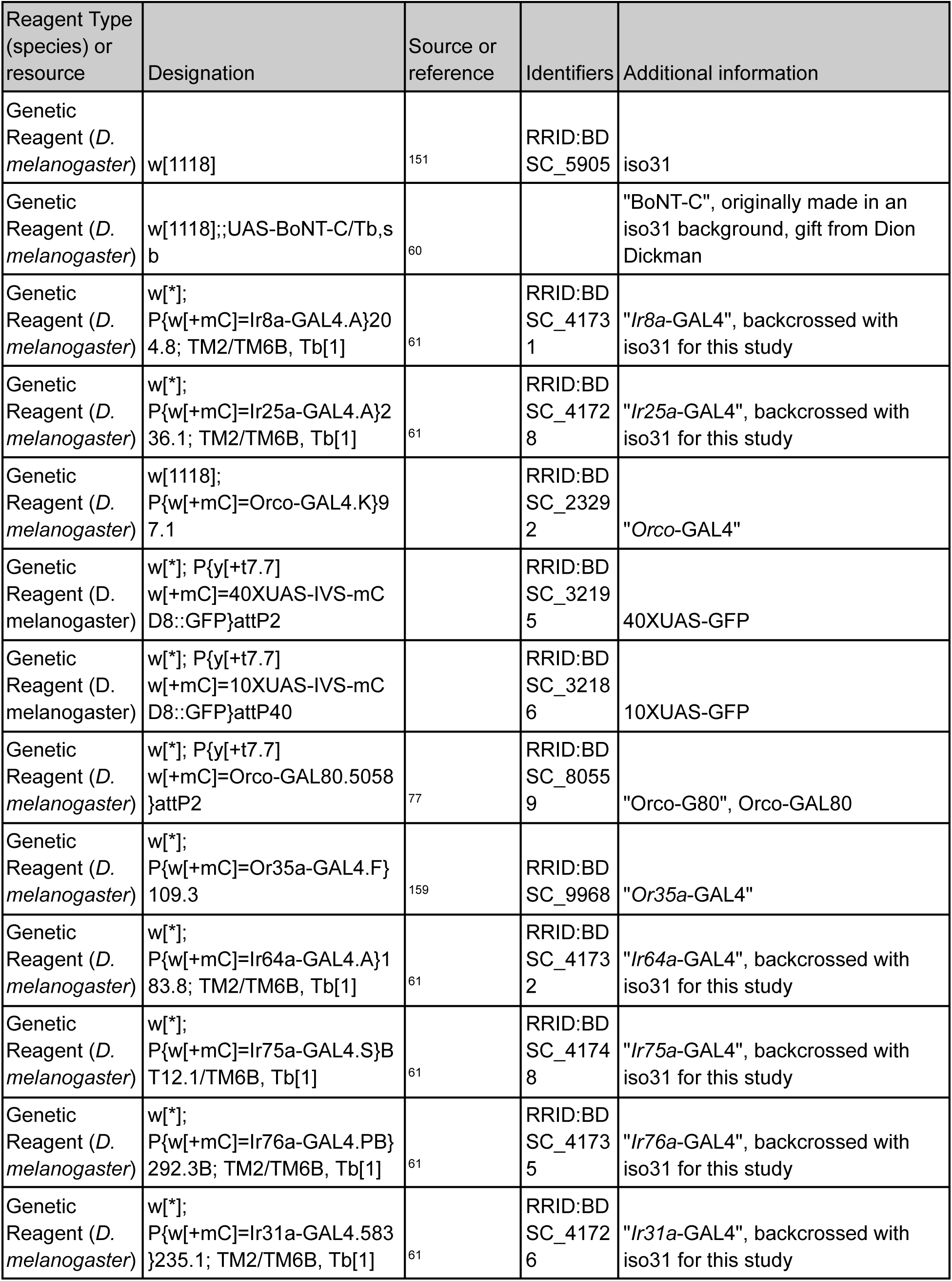
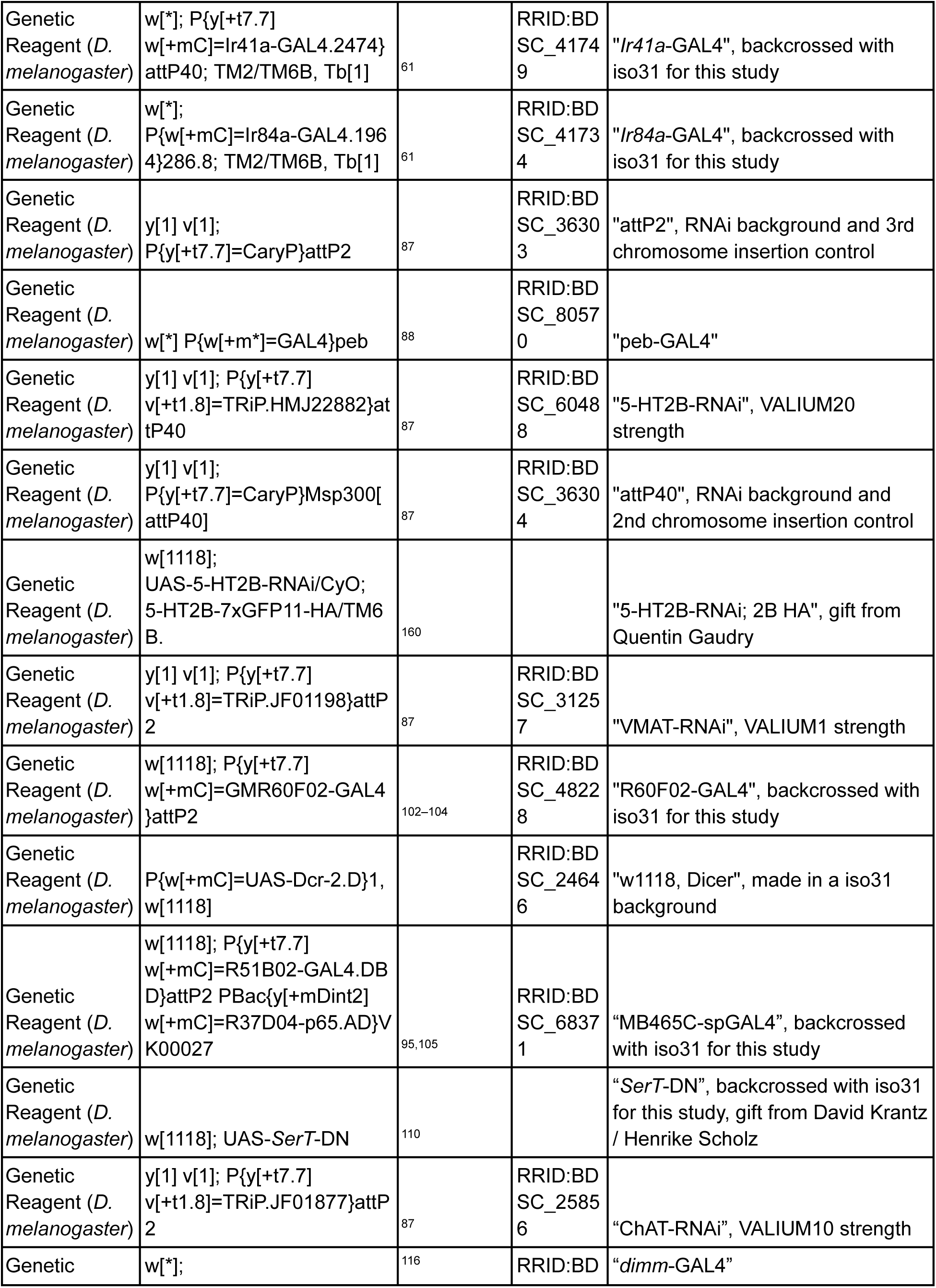
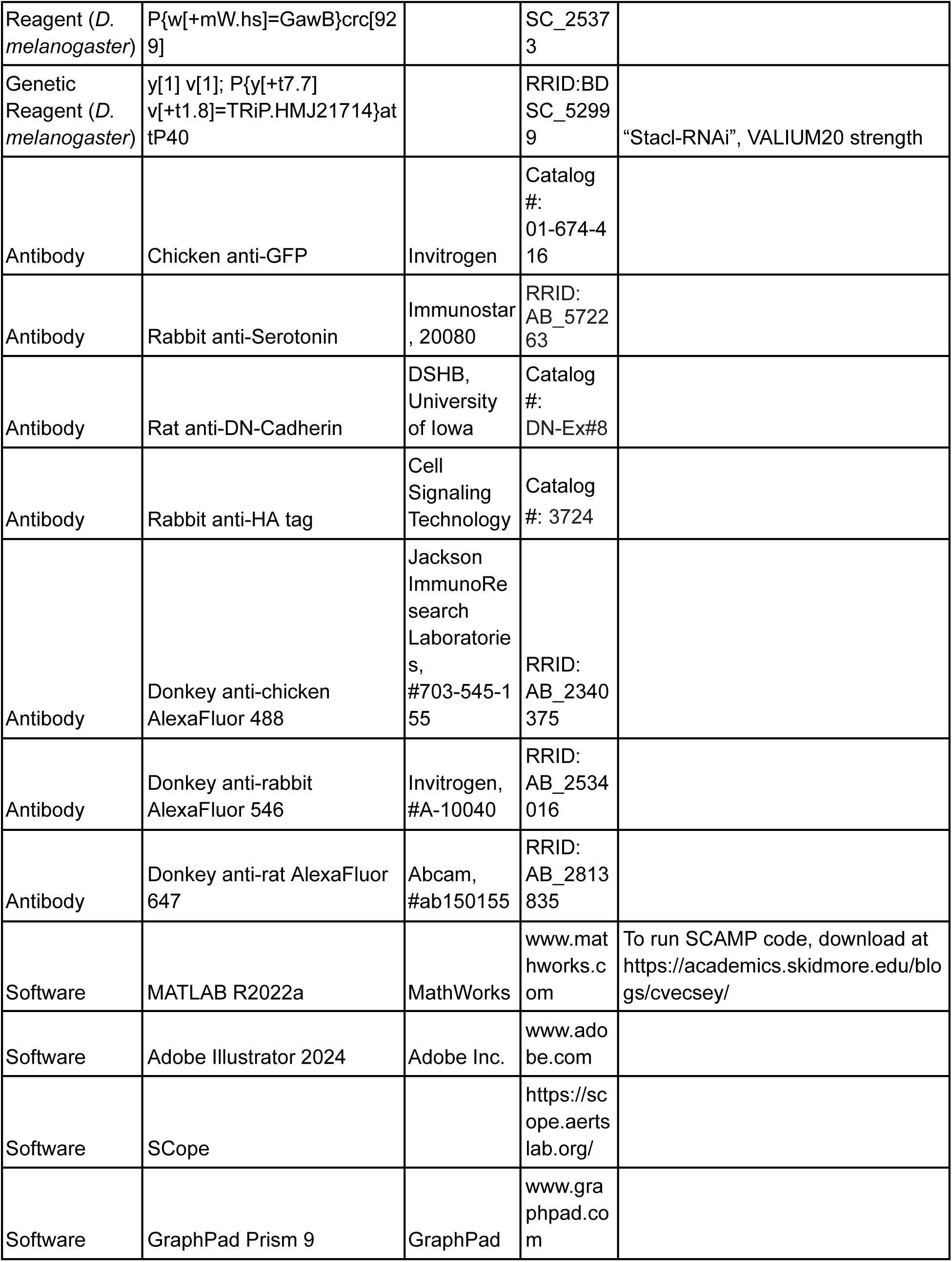
Sources and identifiers of all key reagents and resources.

**Table 2.**
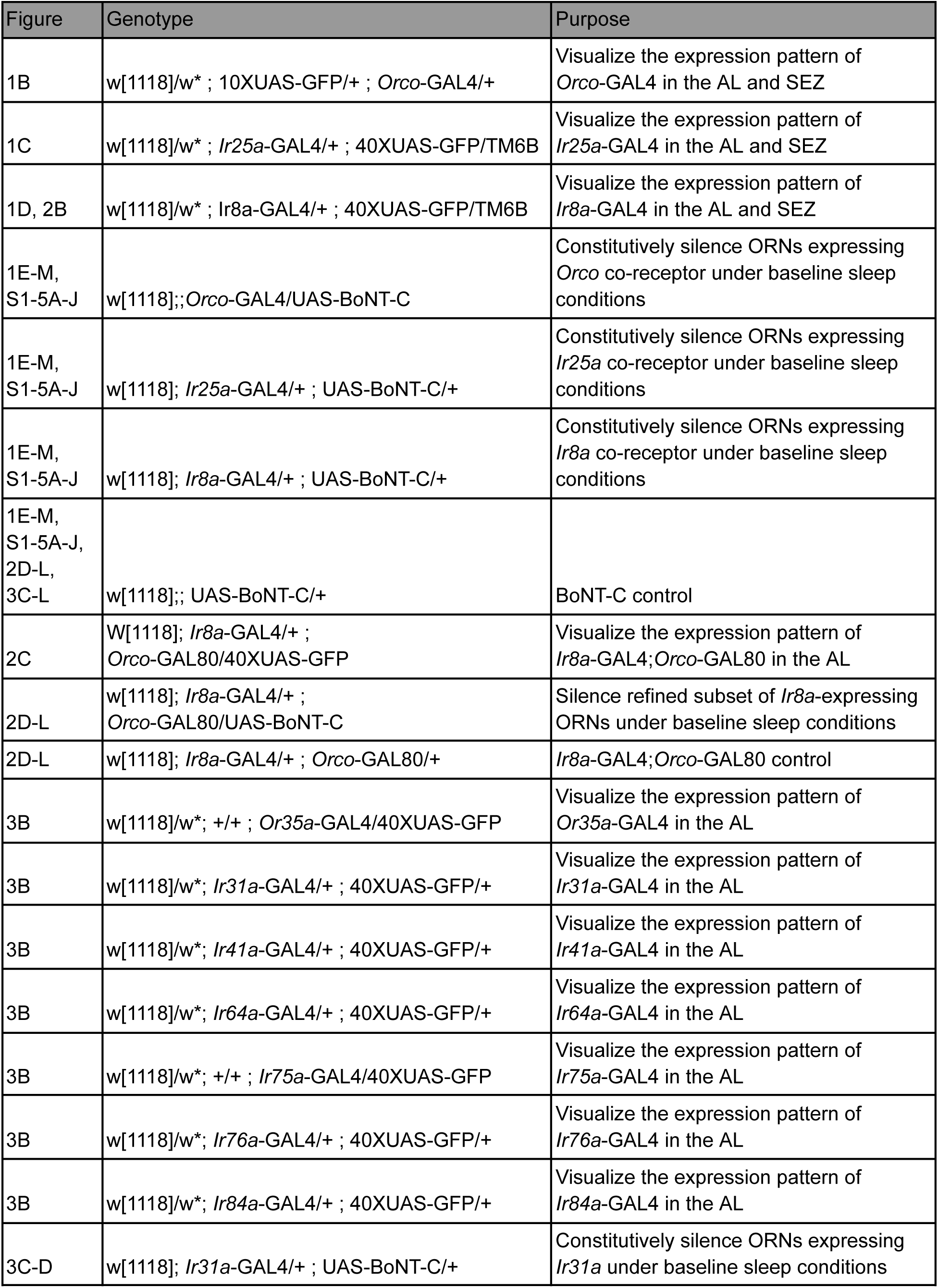
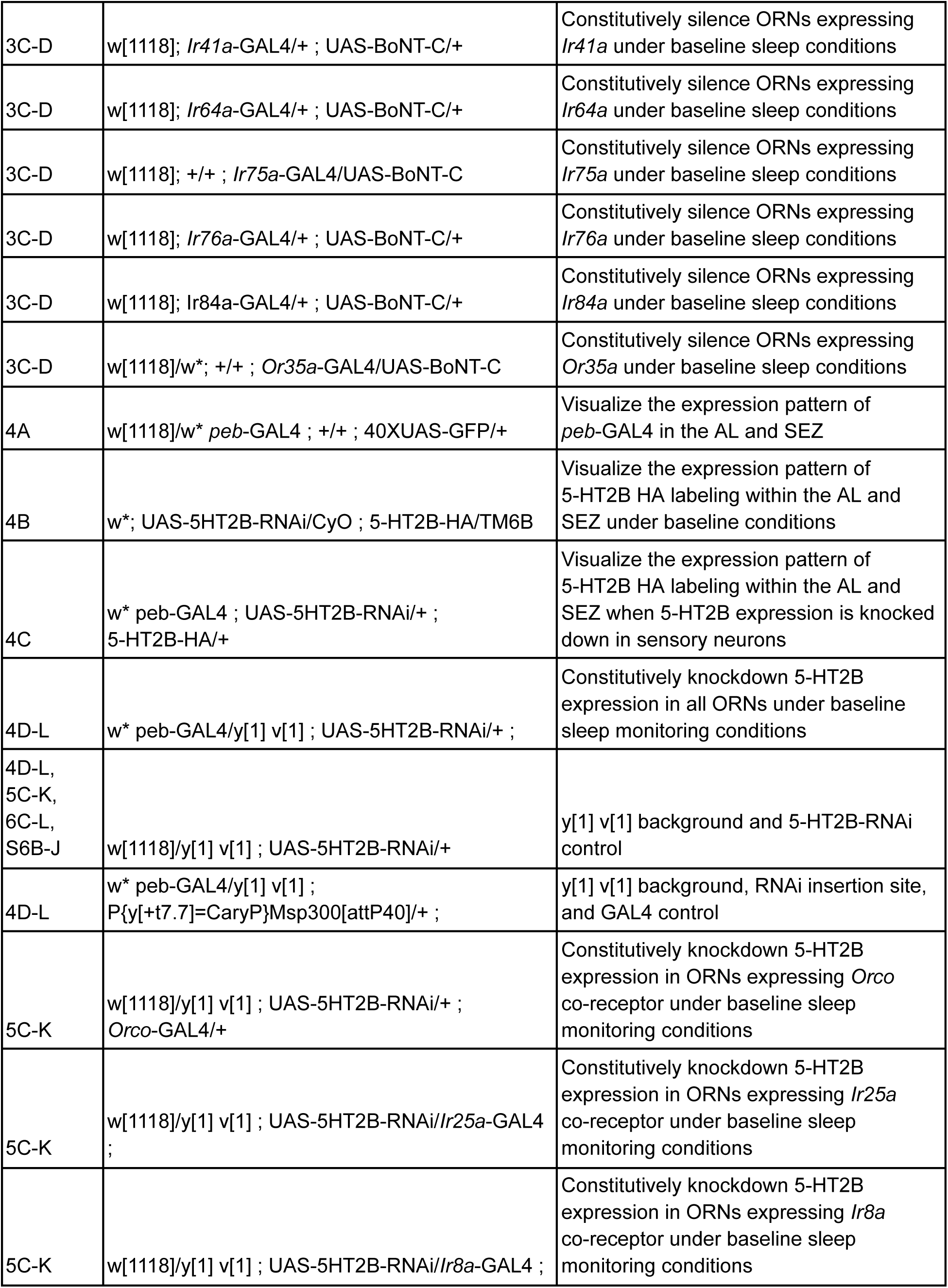
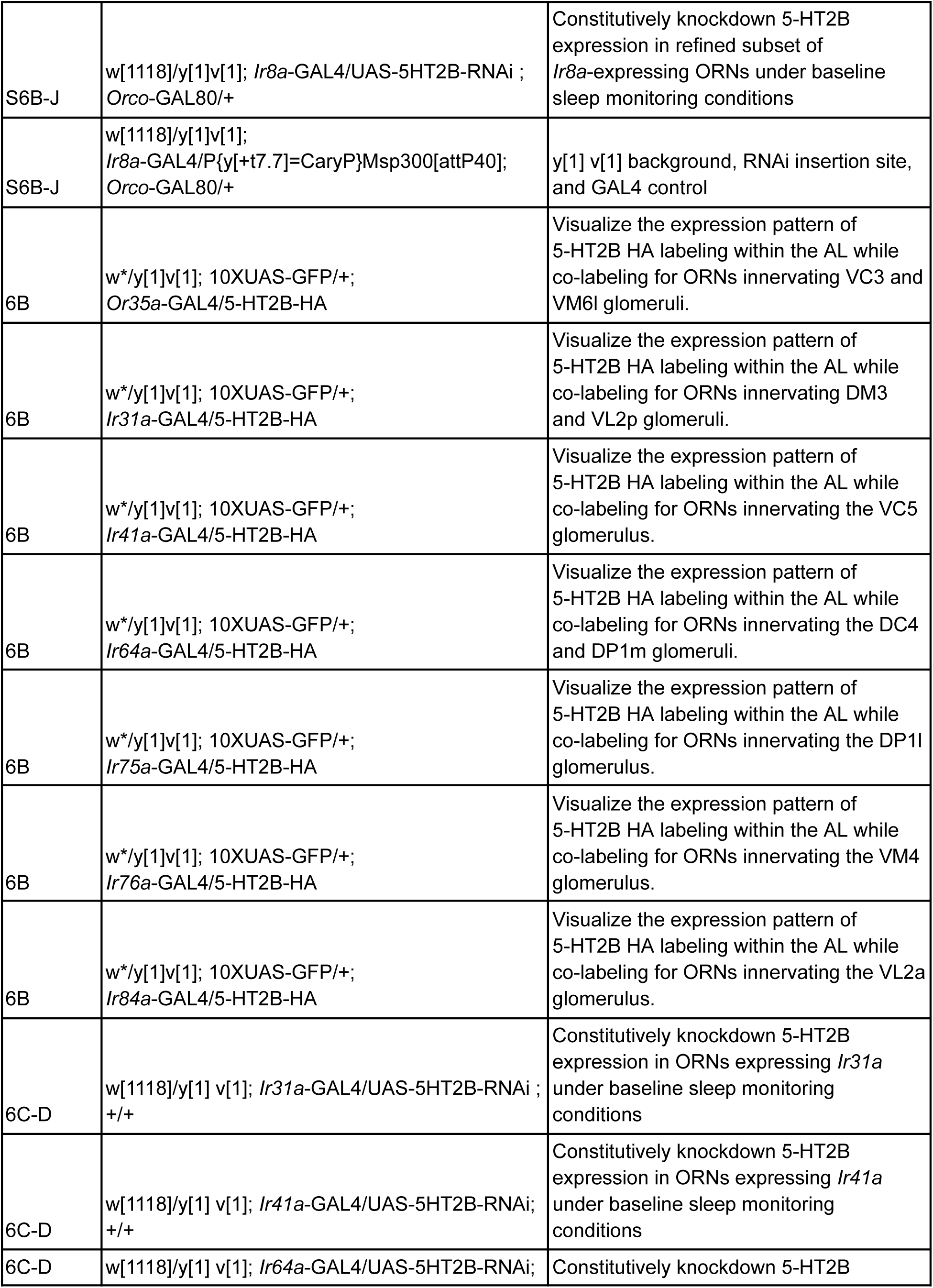
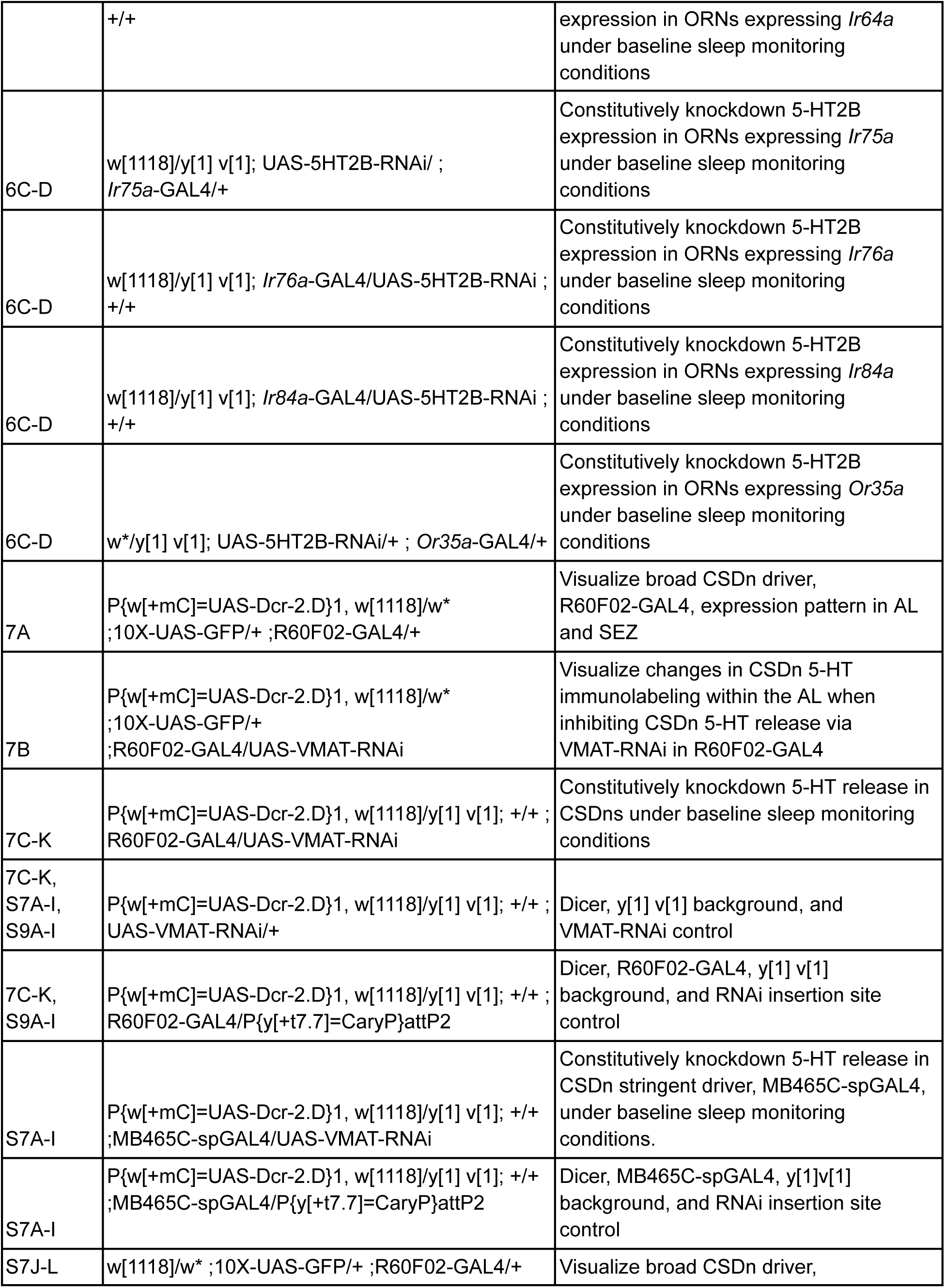
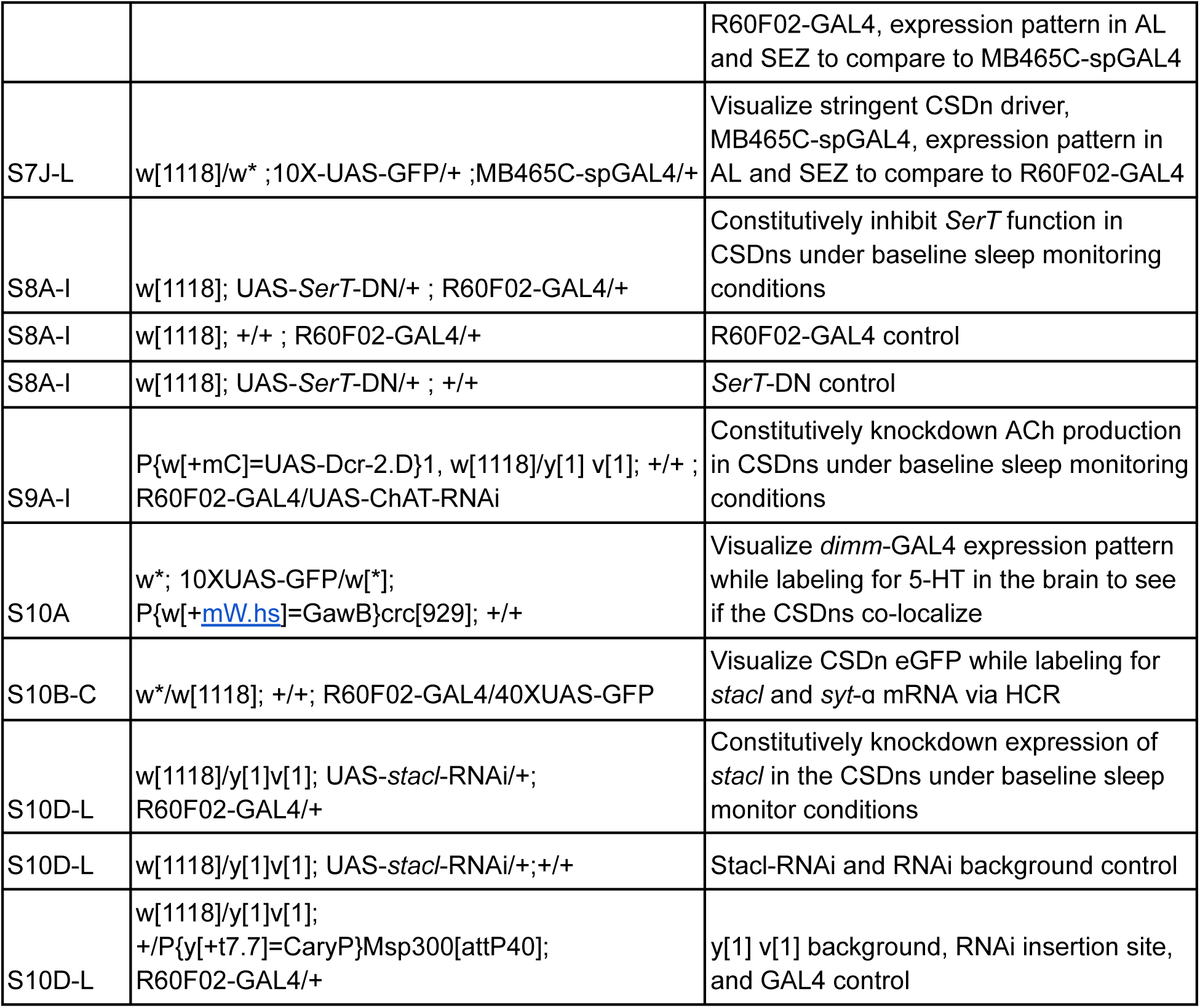
Genotypes of flies used in each figure.

### Drosophila activity monitor (DAM) sleep

Flies were anesthetized with CO_2_ and separated by sex and mating status under a stereoscope into vials of 2% sucrose and 5% agar diet (DAM diet) approximately 2-3 days prior to placement into DAM assays. When ensuring mated flies, genotypes were stored at a 2:1 female:male ratio per vial. When ensuring virgin flies, virgin females were identified by visualizing their meconium and were stored without males. Once 3-7 days of age, flies of each condition per genotype were anesthetized on ice and individually loaded into pyrex glass tubes (Trikinetics, PGT5x65) containing the DAM diet sealed on one end with unscented dental wax and on the other end with a bit of cotton. Tubes with flies were then housed in individual channels of DAM2 monitors (Trikinetics). Activity counts were measured at a 1-minute acquisition frequency across 5 days in 12:12 light/dark cycles. Days 1-2 were treated as acclimation and omitted from analysis. Following the trials, individual fly data were extracted using the DAM File Scan program and channel files were analyzed using the MATLAB software SCAMP^123^. SCAMP was used to determine a variety of standard sleep metrics such as changes in total sleep duration, activity per waking minute, number and duration of sleep episodes, in addition to measures of sleep depth/arousal (pwake) and sleep drive (pdoze)^70^. Data was then pooled across trials and analyzed for statistical significance using GraphPad Prism 9.5.1.

Normally distributed data was tested for significance using a one-way ANOVA and Tukey posttest. In experiments screening multiple GAL4 lines, normally distributed data was instead tested using a Dunnett’s posttest. Otherwise, a nonparametric Kruskall-Wallis test with Dunn’s posttest was used. In all graphs, *p<0.05, **p<0.01, ***p<0.001, ****p<0.0001.

### Immunocytochemistry and image acquisition

Intact brains were dissected at 3-7 days of age in *Drosophila* saline (CSHL recipe) and fixed in 4% paraformaldehyde on ice for 30 minutes-1 hour. Samples were then taken through a series of 4x 15 minute PBST (PBS with 0.5% Triton X-100) washes and blocked for 1 hour in blocking solution (consisting of 2-4% IgG free BSA (Jackson ImmunoResearch, CAS:001-000-162) in PBSAT (PBST with 5mM sodium azide). Samples were then incubated for 48 hours at 4°C with agitation in primary antibody solution (consisting of primary antibodies in blocking solution as above). Next, samples were washed, blocked, and incubated as before except now in secondary antibody solution (secondary antibodies in blocking solution). Lastly, samples were taken through a series of 2x 15 minute PBST, 2x 15 minute PBS, and 1x 10 minute glycerol (40%, 60%, and 80% glycerol in DI water) washes. When labeling for 5-HT2B HA, we adjusted our protocol as such: fixing samples at room temperature, blocking in 10% Normal goat serum (NGS, Thermo Fisher Scientific) and applying antibodies in 5% NGS^85^. Afterward, samples were mounted in VectaShield (Vector Labs Burlingame, CA #H-1000). Samples were scanned using an Olympus confocal microscope FV1000 equipped with 40x silicon oil immersion lens. Images were viewed and analyzed using Olympus FluoView software and positioned using Adobe Illustrator 2024.

### Hybridization Chain Reaction (HCR)

All supplies for HCR were purchased from Molecular Instruments Inc. (Los Angeles, CA). Brains were dissected in PBS and fixed in 4% paraformaldehyde for 30 minutes at RT. Brains were then placed in a series of 15-minute PBST washes as mentioned previously. Brains were then transferred to hybridization buffer for 30 minutes at 37℃. Afterwards, probe solution (4µl of each probe set per 1000µl of hybridization buffer) was applied and brains were incubated overnight at 37℃. The following day, brains were washed with probe wash buffer in 15-minute intervals at 37℃. Brains were then placed in a 5-minute wash series with 5x SSCT at RT. Afterwards, brains were transferred to amplification buffer for 30 minutes at RT. While being pre-amplified, 10µl aliquots of 3µM hairpin stock solutions were snap-cooled in a thermocycler (Bio-Rad T100 Thermal Cycler). The aliquots were heated for 90 seconds at 95℃ and then transferred to a dark drawer for 30 minutes. A hairpin solution was then prepared by adding the snap cooled aliquots (10µl per 500µl of needed solution) to the amplification buffer. Hairpin solution was then applied to brains and they were incubated overnight in the dark at RT. On the third and final day, the brains were washed (2x 5 minute, 2x 30 minute, 2x 5 minute) with 5x SSCT at RT followed by a series of 10-minute glycerol washes (40%, 60%, 80%). The brains were then mounted and scanned as described above.

### Scope

ORN transcriptomic data was queried and downloaded from the publicly available SCope tool (https://scope.aertslab.org). The ORN clusters were identified and selected within the antenna dataset (stringent version) as previously described^93^. All data was exported from the HVG t-SNE coordinate set, log-transformed and counts per million normalized. Expression data from individual ORN types was organized using marker gene criteria where neurons with expression > 0 CPM for a given OR/IR and their corresponding co-receptor from the literature. For example, expression of a given gene from DA1 *Or67d+* neurons would only be identified in cells having both > 0 CPM *Or67d* and *Orco* expression. It is important to note that this approach was taken prior to the publication of the expanded co-receptor atlas that demonstrated many ORN types expressed multiple co-receptors ^46^. However, the reductionist approach we took here would have still included the ORNs of a given type expressing additional co-receptors.

## Author Contributions

Designed research (OMC, KEC, AMD), conducted research (OMC, BTP, GP, SR, IAA), analyzed data (OMC, BTP, GP, IAA), wrote the paper (OMC, AMD).

---

**Supplemental Figure 1.**
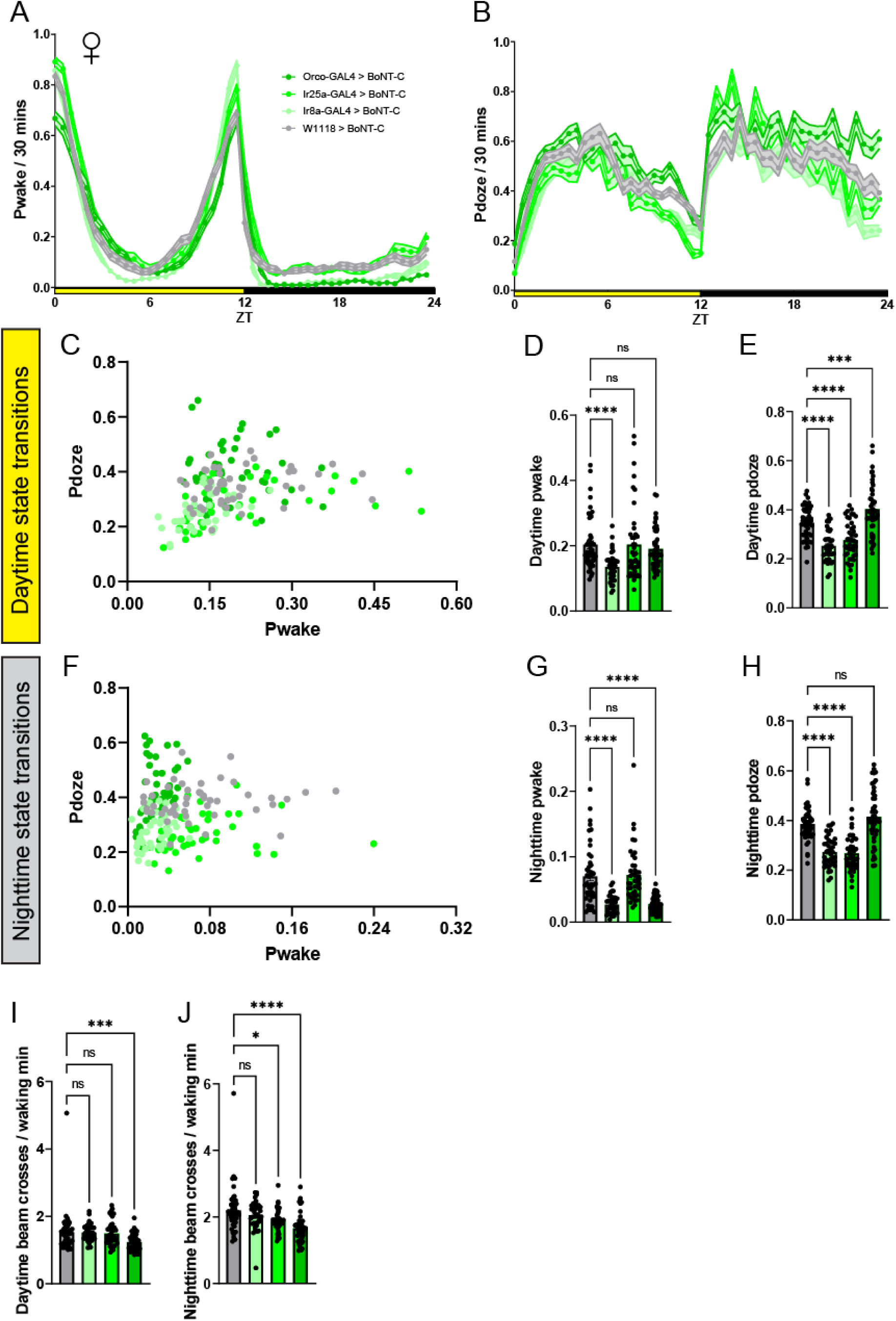
M**a**ted **female ORN co-receptor BoNT-C screen pwake and pdoze architecture. A**) Pwake and **B**) Pdoze per 30 minutes of flies where different ORN co-receptor populations are silenced via BoNT-C. Yellow bar represents daytime (ZT 0-12), while black is nighttime (ZT 12-24). **C**) Scatter plot of the per-fly daytime state transition architecture (comparison of daytime pwake and pdoze) across each tested genotype. **D**) Average daytime pwake. **E**) Average daytime pdoze. **F**) Scatter plot of the per-fly nighttime state transition architecture (comparison of nighttime pwake and pdoze) across each tested genotype. **G**) Average nighttime pwake. **H**) Average nighttime pdoze. **I-J**) Average beam crosses per waking minute in **I**) daytime and **J**) nighttime. In each behavior experiment, 3-7 day age mated female flies were tested, N=39-47 per genotype. Gray = BoNT-C controls, lightest green = *Ir8a*-GAL4 > BoNT-C, brighter green = *Ir25a*-GAL4 > BoNT-C, darkest green = *Orco*-GAL4 > BoNT-C. See methods for details on statistical analysis. *p<0.05, **p<0.01, ***p<0.001, ****p<0.0001.

**Supplemental Figure 2.**
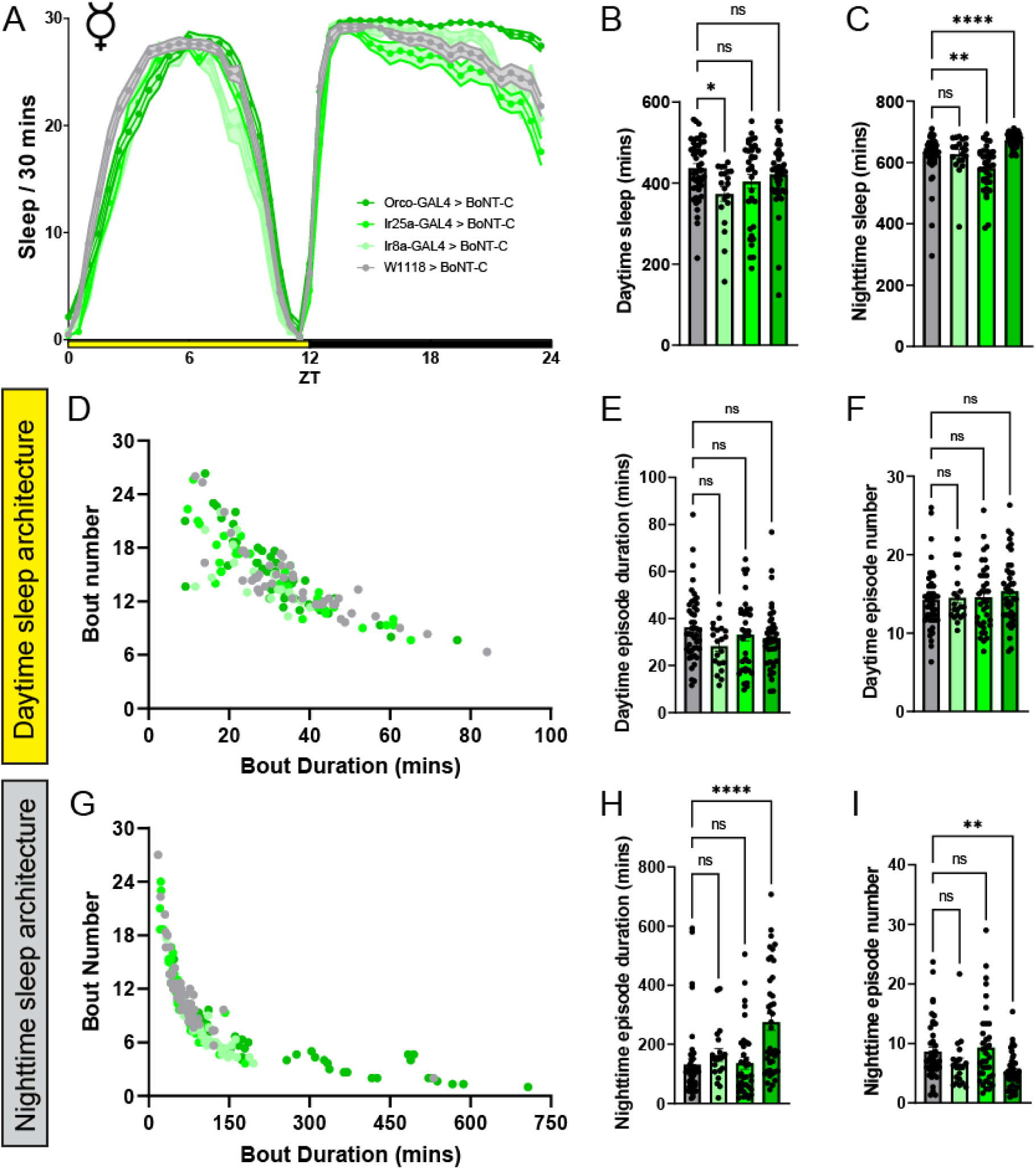
V**i**rgin **female ORN co-receptor BoNT-C screen sleep architecture. A)** Sleep per 30 minutes of flies where different ORN co-receptor populations are silenced via BoNT-C. Yellow bar represents daytime (ZT 0-12), while black is nighttime (ZT 12-24). **B-C**) Average sleep duration of **B**) daytime and **C**) nighttime sleep phases. **D**) Scatter plot of the per-fly daytime sleep architecture across each tested genotype. **E**) Average daytime sleep bout duration. **F**) Average daytime sleep bout number. **G**) Scatter plot of the per-fly nighttime sleep architecture. **H**) Average nighttime sleep bout duration. **I**) Average nighttime sleep bout number. In each behavior experiment, 3-7 day age virgin female flies were tested, N=20-45 per genotype. Gray = BoNT-C controls, lightest green = *Ir8a*-GAL4 > BoNT-C, brighter green = *Ir25a*-GAL4 > BoNT-C, darkest green = *Orco*-GAL4 > BoNT-C. See methods for details on statistical analysis. *p<0.05, **p<0.01, ***p<0.001, ****p<0.0001.

**Supplemental Figure 3.**
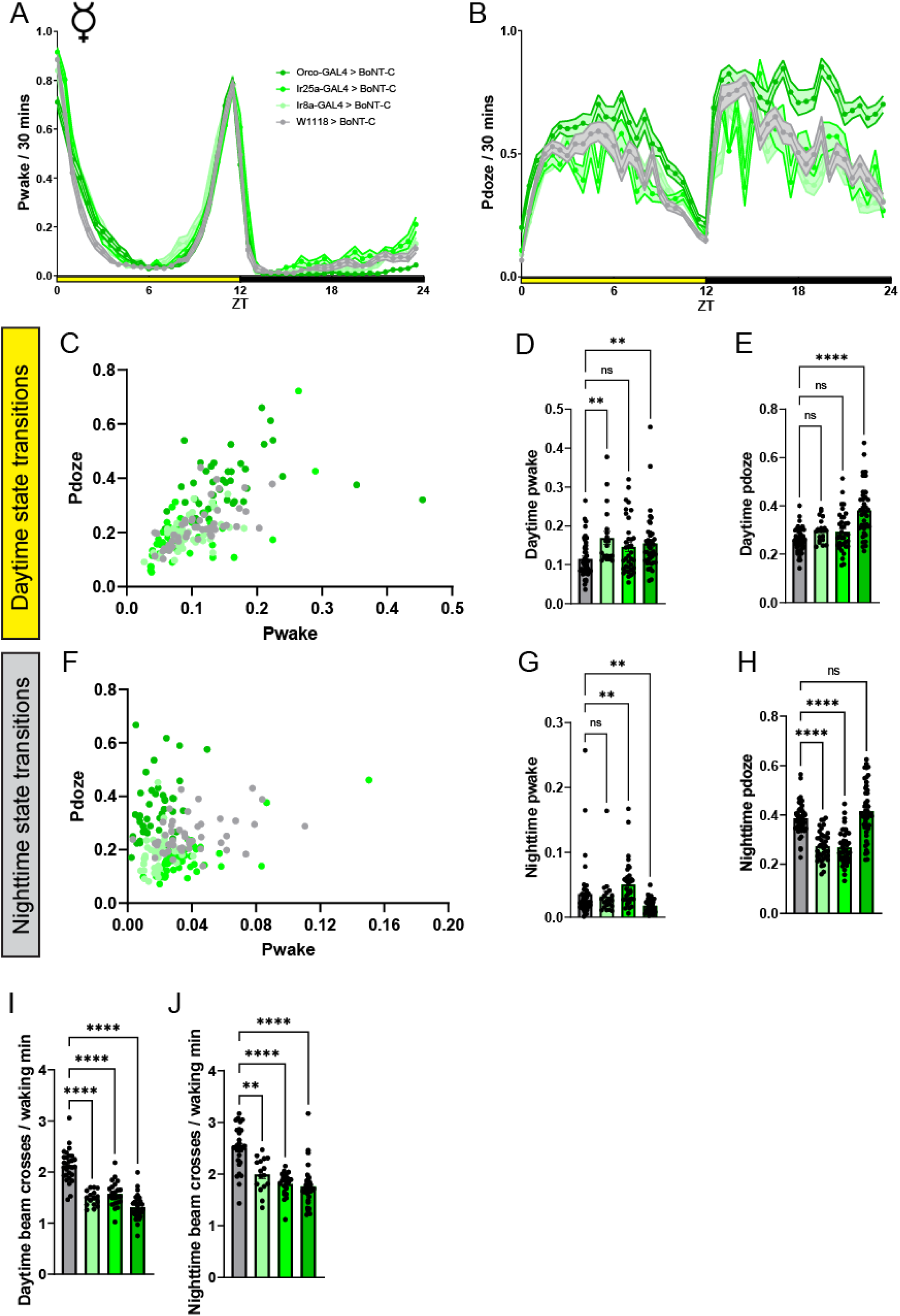
V**i**rgin **female ORN co-receptor BoNT-C screen pwake and pdoze architecture. A**) Pwake and **B**) Pdoze per 30 minutes of flies where different ORN co-receptor populations are silenced via BoNT-C. Yellow bar represents daytime (ZT 0-12), while black is nighttime (ZT 12-24). **C**) Scatter plot of the per-fly daytime state transition architecture across each tested genotype. **D**) Average daytime pwake. **E**) Average daytime pdoze. **F**) Scatter plot of the per-fly nighttime state transition architecture across each tested genotype. **G**) Average nighttime pwake. **H**) Average nighttime pdoze. **I-J**) Average beam crosses per waking minute in **I)** daytime and **J**) nighttime. In each behavior experiment, 3-7 day age virgin female flies were tested, N=20-45 per genotype. Gray = BoNT-C controls, lightest green = *Ir8a*-GAL4 > BoNT-C, brighter green = *Ir25a*-GAL4 > BoNT-C, darkest green = *Orco*-GAL4 > BoNT-C. See methods for details on statistical analysis. *p<0.05, **p<0.01, ***p<0.001, ****p<0.0001.

**Supplemental Figure 4.**
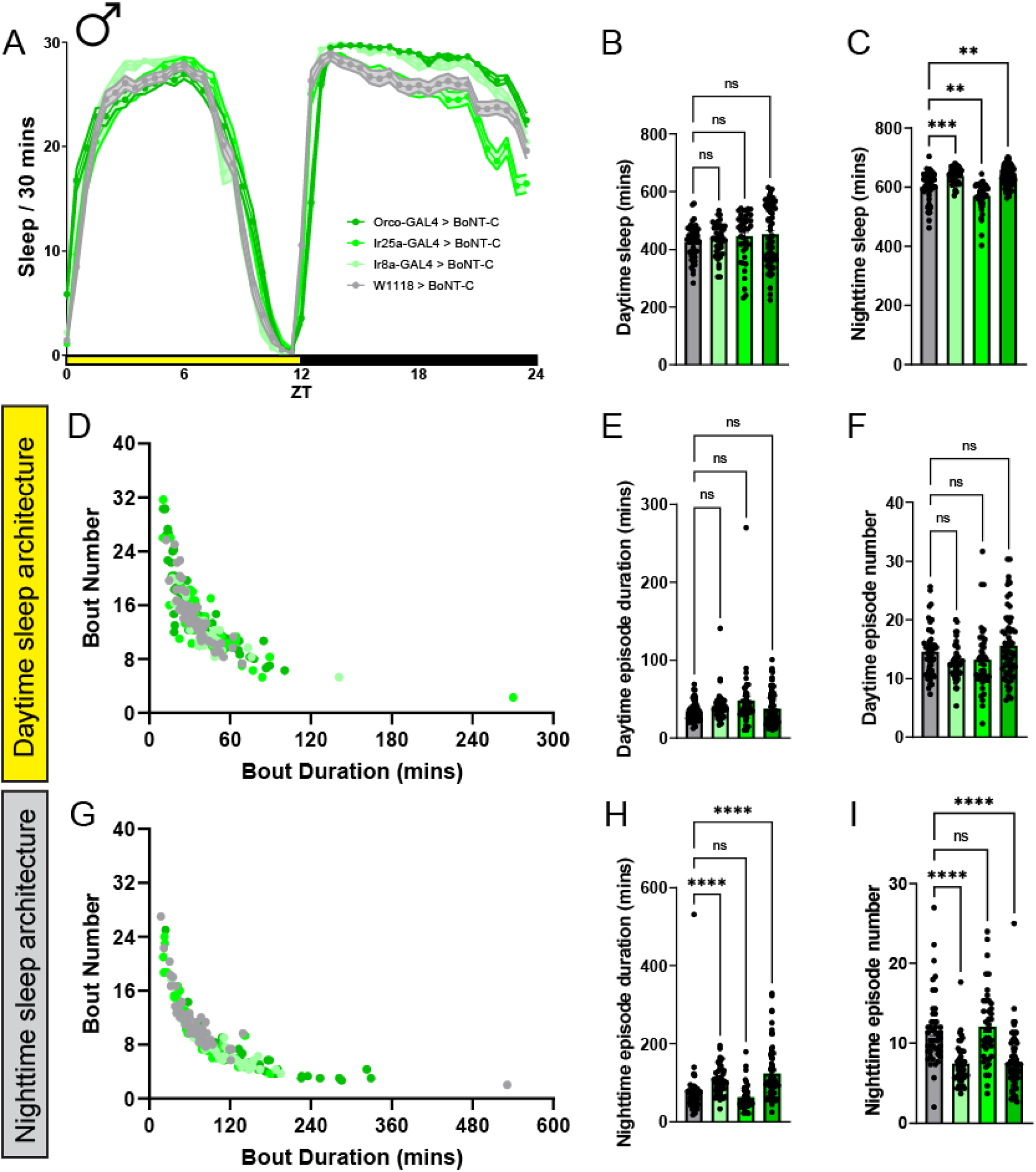
M**a**ted **male ORN co-receptor BoNT-C screen sleep architecture. A)** Sleep per 30 minutes of flies where different ORN co-receptor populations are silenced via BoNT-C. Yellow bar represents daytime (ZT 0-12), while black is nighttime (ZT 12-24). **B-C**) Average sleep duration of **B**) daytime and **C**) nighttime sleep phases. **D**) Scatter plot of the per-fly daytime sleep architecture across each tested genotype. **E**) Average daytime sleep bout duration. **F**) Average daytime sleep bout number. **G**) Scatter plot of the per-fly nighttime sleep architecture across each genotype. **H**) Average nighttime sleep bout duration. **I**) Average nighttime sleep bout number. In each behavior experiment, 3-7 day age mated male flies were tested, N=41-64 per genotype. Gray = BoNT-C controls, lightest green = *Ir8a*-GAL4 > BoNT-C, brighter green = *Ir25a*-GAL4 > BoNT-C, darkest green = *Orco*-GAL4 > BoNT-C. See methods for details on statistical analysis. *p<0.05, **p<0.01, ***p<0.001, ****p<0.0001.

**Supplemental Figure 5.**
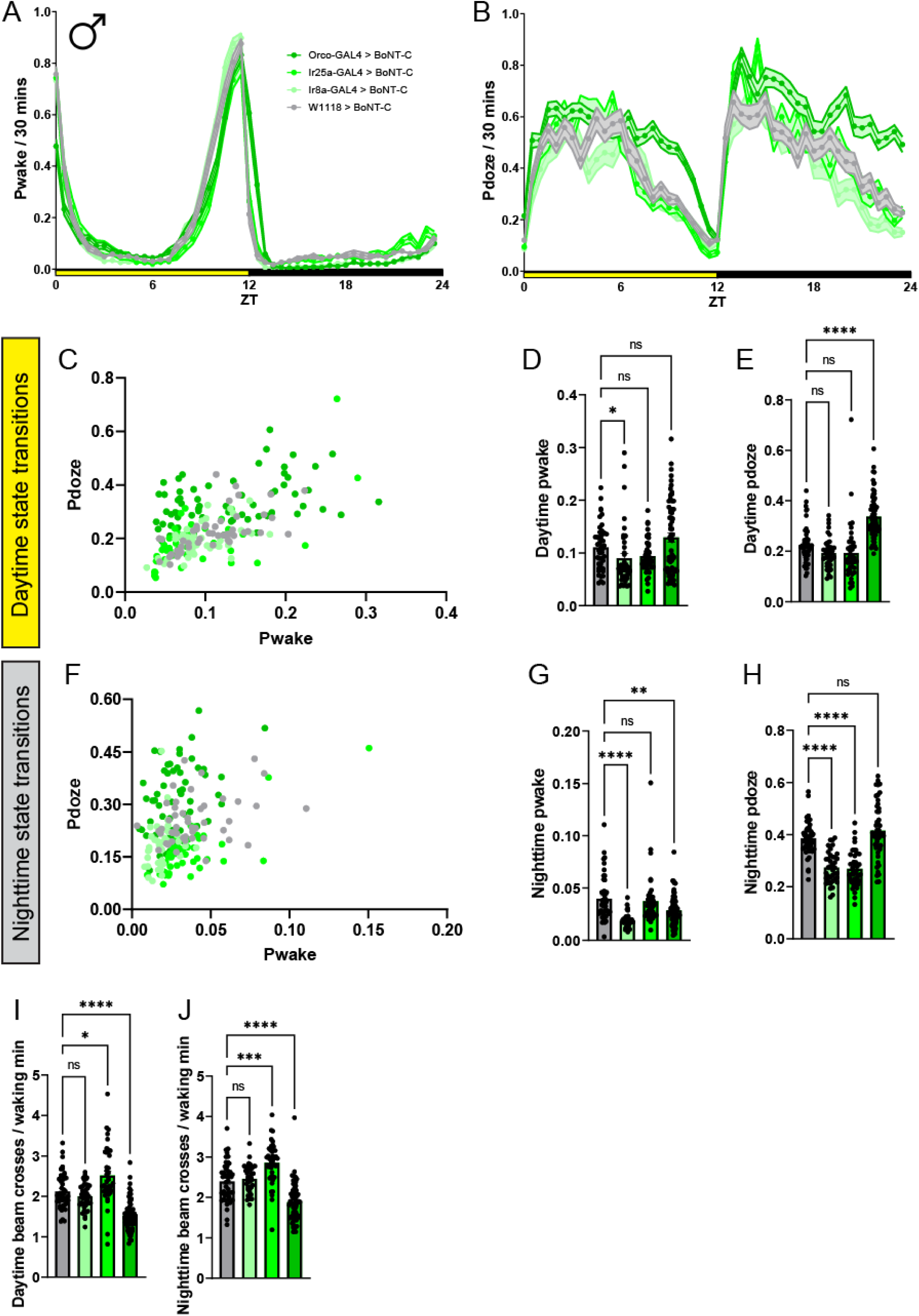
M**a**ted **male ORN co-receptor BoNT-C screen pwake and pdoze architecture.** Pwake and **B**) Pdoze per 30 minutes of flies where different ORN co-receptor populations are silenced via BoNT-C. Yellow bar represents daytime (ZT 0-12), while black is nighttime (ZT 12-24). **C**) Scatter plot of the per-fly daytime state transition architecture across each tested genotype. **D**) Average daytime pwake. **E**) Average daytime pdoze. **F**) Scatter plot of the per-fly nighttime state transition architecture across each tested genotype. **G**) Average nighttime pwake. **H**) Average nighttime pdoze. **I-J**) Average beam crosses per waking minute in **I**) daytime and **J**) nighttime. In each behavior experiment, 3-7 day age mated male flies were tested, N=41-64 per genotype. Gray = BoNT-C controls, lightest green = *Ir8a*-GAL4 > BoNT-C, brighter green = *Ir25a*-GAL4 > BoNT-C, darkest green = *Orco*-GAL4 > BoNT-C. See methods for details on statistical analysis. *p<0.05, **p<0.01, ***p<0.001, ****p<0.0001.

**Supplemental Figure 6.**
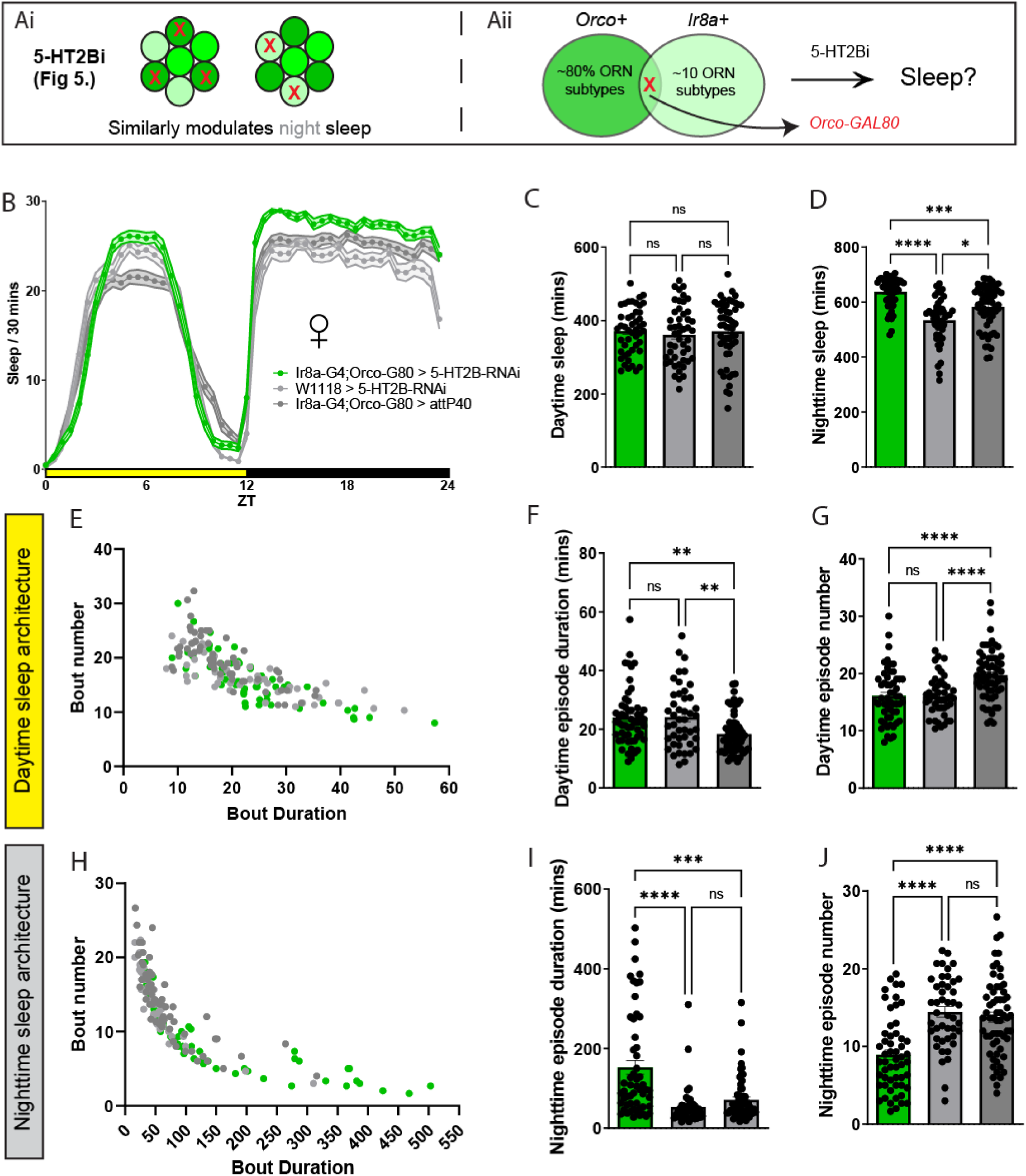
A**l**tering **the expression of *Ir8a*-GAL4 does not reduce the effect of 5-HT2B-RNAi on nighttime sleep. A)** Schematic of hypothesis testing the contributions 5-HT2B expression of individual ORN types on sleep. Since knocking down 5-HT2B expression in both the *Orco* and *Ir8a*-GAL4 drivers similarly influenced nighttime sleep architecture (**Figure 5**), we questioned whether knocking down 5-HT2B within ORNs labeled in an altered expression pattern of *Ir8a*-GAL4 via an *Orco*-GAL80 would similarly modulate nighttime sleep. **B**) Sleep per 30 minutes of flies where 5-HT2B expression is knocked down in ORNs labeled in by an altered expression pattern of *Ir8a*-GAL4. Yellow bar represents daytime (ZT 0-12), while black is nighttime (ZT 12-24). **C-D**) Average sleep duration of **C**) daytime and **D**) nighttime sleep phases. **E**) Scatter plot of the per-fly daytime sleep architecture across each tested genotype. **F**) Average daytime sleep bout duration. **G**) Average daytime sleep bout number. **H**) Scatter plot of the per-fly nighttime sleep architecture. **I**) Average nighttime sleep bout duration. **J**) Average nighttime sleep bout number. In each behavior experiment, 3-7 day age mated female flies were tested, N=45-61 per genotype. Lightest gray = 5-HT2B-RNAi control, darker gray = Ir8a-GAL4;Orco-GAL80 control, green = *Ir8a*-GAL4;*Orco*-GAL80 > 5-HT2B-RNAi.

**Supplemental Figure 7.**
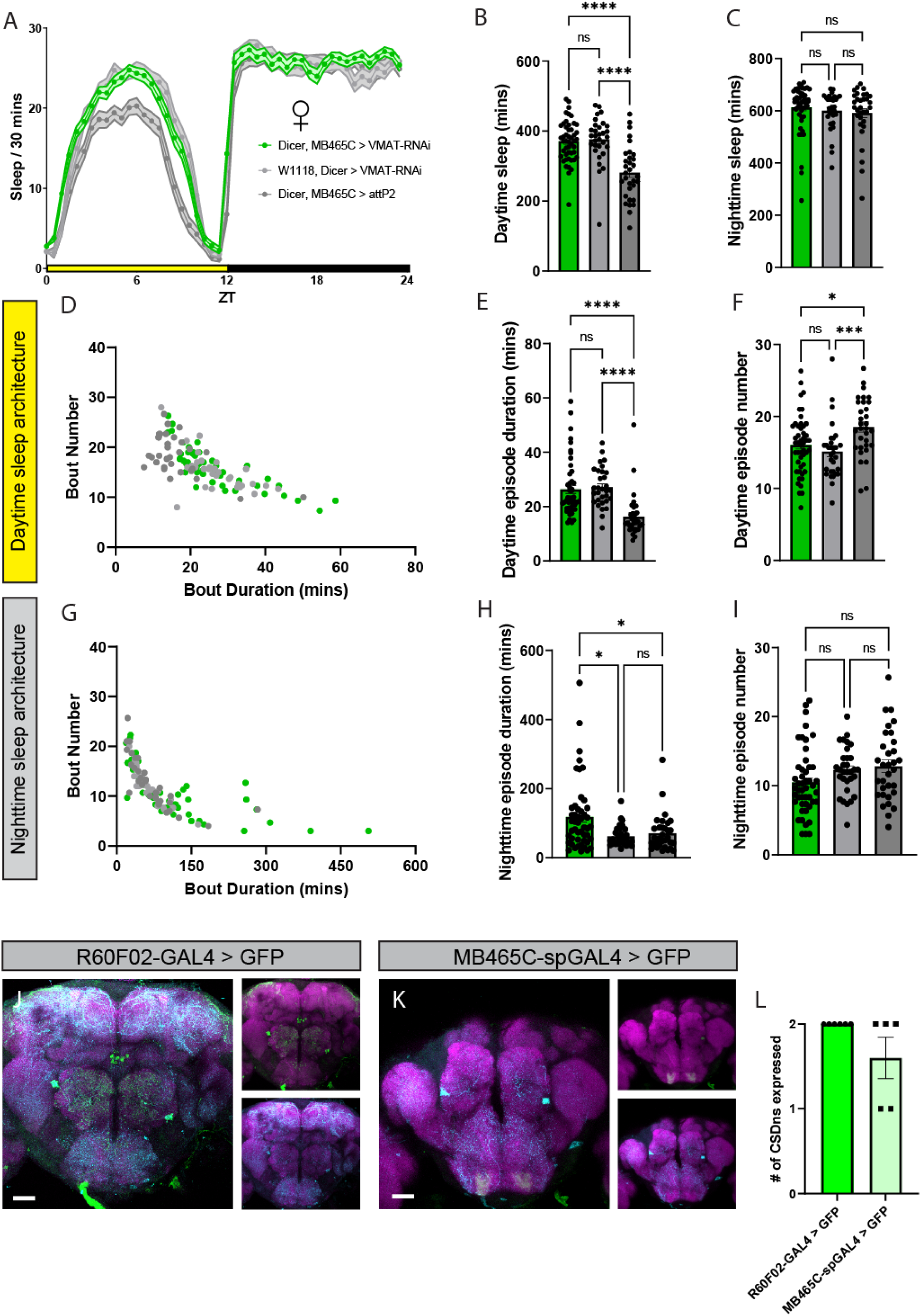
E**f**fects **of VMAT-RNAi in a stringent CSDn driver line. A**) Sleep per 30 minutes of flies where CSDn 5-HT signaling is inhibited via Dicer, MB465C-spGAL4 > VMAT-RNAi. Yellow bar represents daytime (ZT 0-12), while black is nighttime (ZT 12-24). **B-C**) Average sleep duration of **B**) daytime and **C**) nighttime sleep phases. **D**) Scatter plot of the per-fly daytime sleep architecture across each tested genotype. **E**) Average daytime sleep bout duration. **F**) Average daytime sleep bout number. **G**) Scatter plot of the per-fly nighttime sleep architecture across each genotype. **H**) Average nighttime sleep bout duration. **I**) Average nighttime sleep bout number. In each behavior experiment, 3-7 day age mated male flies were tested, N=31-48 per genotype. Lightest gray = Dicer and VMAT-RNAi control, darker gray = Dicer, R60F02-GAL4 control, green = Dicer, R60F02-GAL4 > VMAT-RNAi. **J-K**) Comparison of glomerular innervation patterns of CSDns driving GFP signal in **A**) R60F02-GAL4 and **B**) MB465C-spGAL4. GFP signal driven in the GAL4 (green), 5-HT (cyan), and the neuropil is labeled by N-cadherin (magenta). Scale bar is 30µm. **L**) Quantification of the number of CSDns labeled per brain by either GAL4 (2 CSDns max per brain). N=5-6 brains per genotype. See methods for details on statistical analysis. *p<0.05, **p<0.01, ***p<0.001, ****p<0.0001.

**Supplemental Figure 8.**
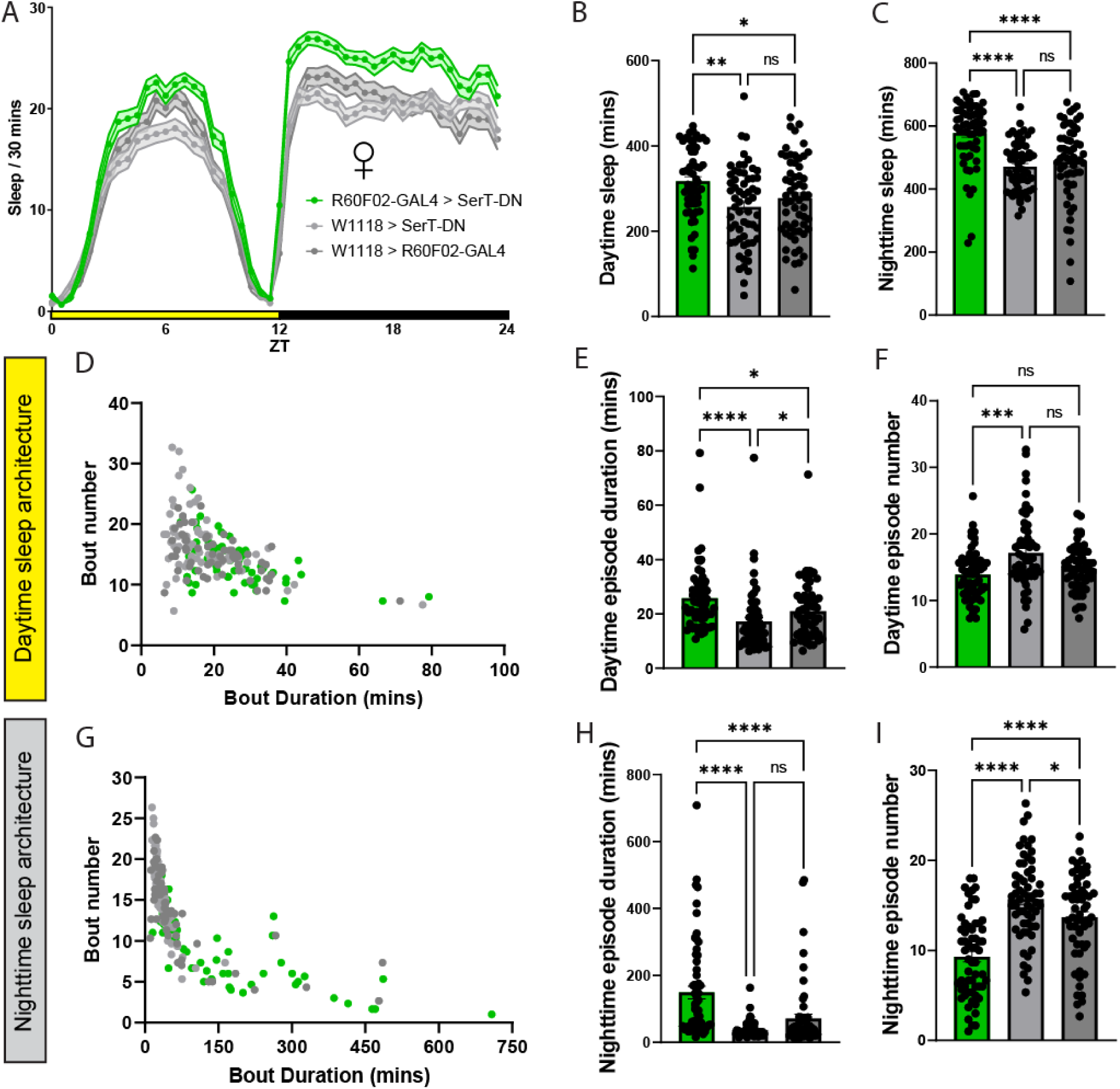
C**S**Dn ***SerT* signaling predominantly modulates nighttime sleep architecture. A**) Sleep per 30 minutes of flies where *SerT* signaling is inhibited in the CSDns via R60F02-GAL4 > *SerT*-DN. Yellow bar represents daytime (ZT 0-12), while black is nighttime (ZT 12-24). **B-C**) Average sleep duration of **B**) daytime and **C**) nighttime sleep phases. **D**) Scatter plot of the per-fly daytime sleep architecture across each tested genotype. **E**) Average daytime sleep bout duration. **F**) Average daytime sleep bout number. **G**) Scatter plot of the per-fly nighttime sleep architecture across each genotype. **H**) Average nighttime sleep bout duration. **I**) Average nighttime sleep bout number. In each behavior experiment, 3-7 day age mated male flies were tested, N=59-60 per genotype. Lightest gray = *SerT*-DN control, darker gray = R60F02-GAL4 control, green = R60F02-GAL4 > *SerT*-DN.

**Supplemental Figure 9.**
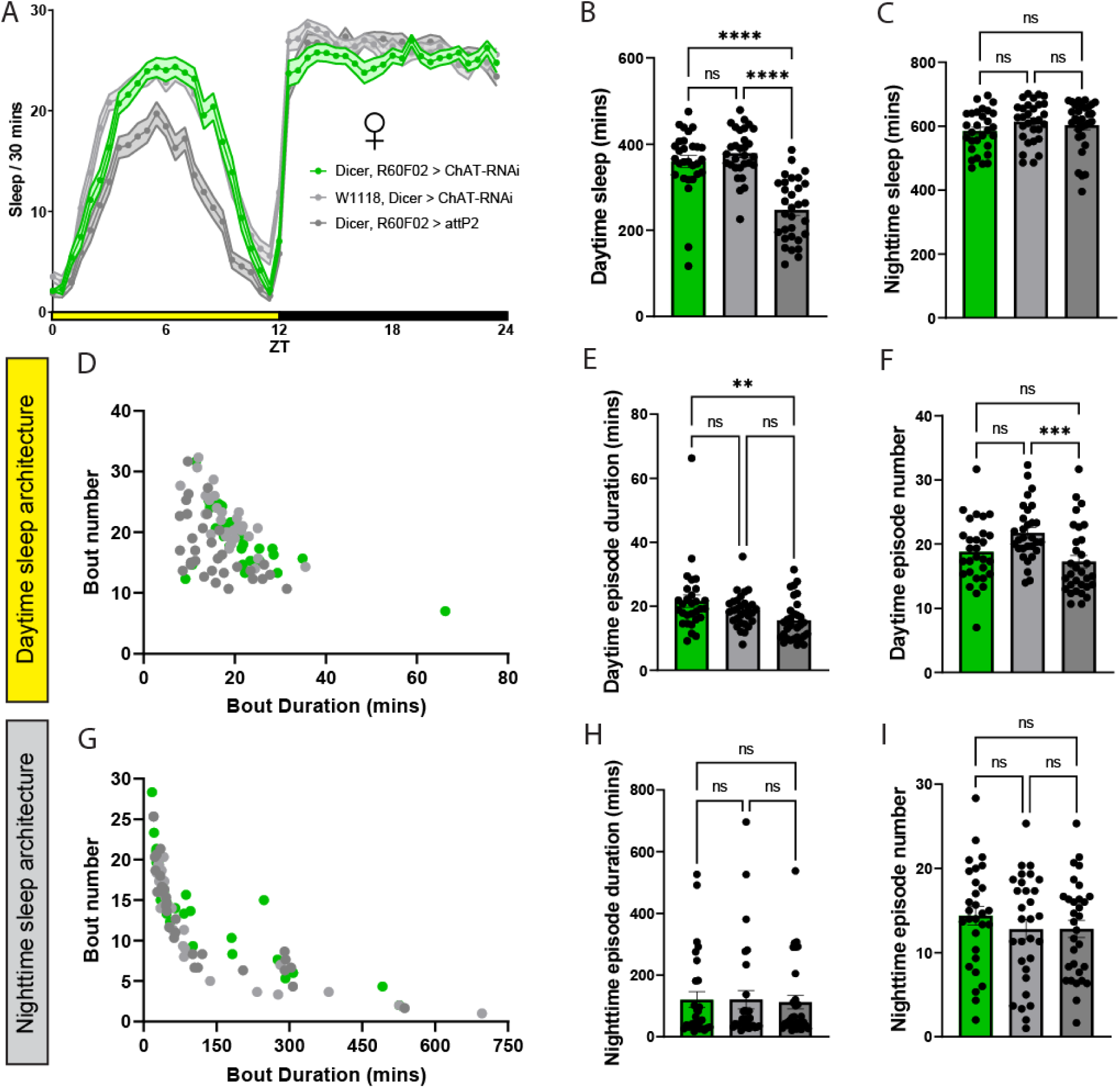
C**S**Dn **ACh signaling does not modulate sleep architecture. A**) Sleep per 30 minutes of flies where ACh signaling is inhibited in the CSDns via Dicer, R60F02-GAL4 > ChAT-RNAi. Yellow bar represents daytime (ZT 0-12), while black is nighttime (ZT 12-24). **B-C**) Average sleep duration of **B**) daytime and **C**) nighttime sleep phases. **D**) Scatter plot of the per-fly daytime sleep architecture across each tested genotype. **E**) Average daytime sleep bout duration. **F**) Average daytime sleep bout number. **G**) Scatter plot of the per-fly nighttime sleep architecture across each genotype. **H**) Average nighttime sleep bout duration. **I**) Average nighttime sleep bout number. In each behavior experiment, 3-7 day age mated male flies were tested, N=29-31 per genotype. Lightest gray = Dicer and ChAT-RNAi control, darker gray = Dicer and R60F02-GAL4 control, green = Dicer, R60F02-GAL4 > ChAT-RNAi. See methods for details on statistical analysis. *p<0.05, **p<0.01, ***p<0.001, ****p<0.0001.

**Supplemental Figure 10.**
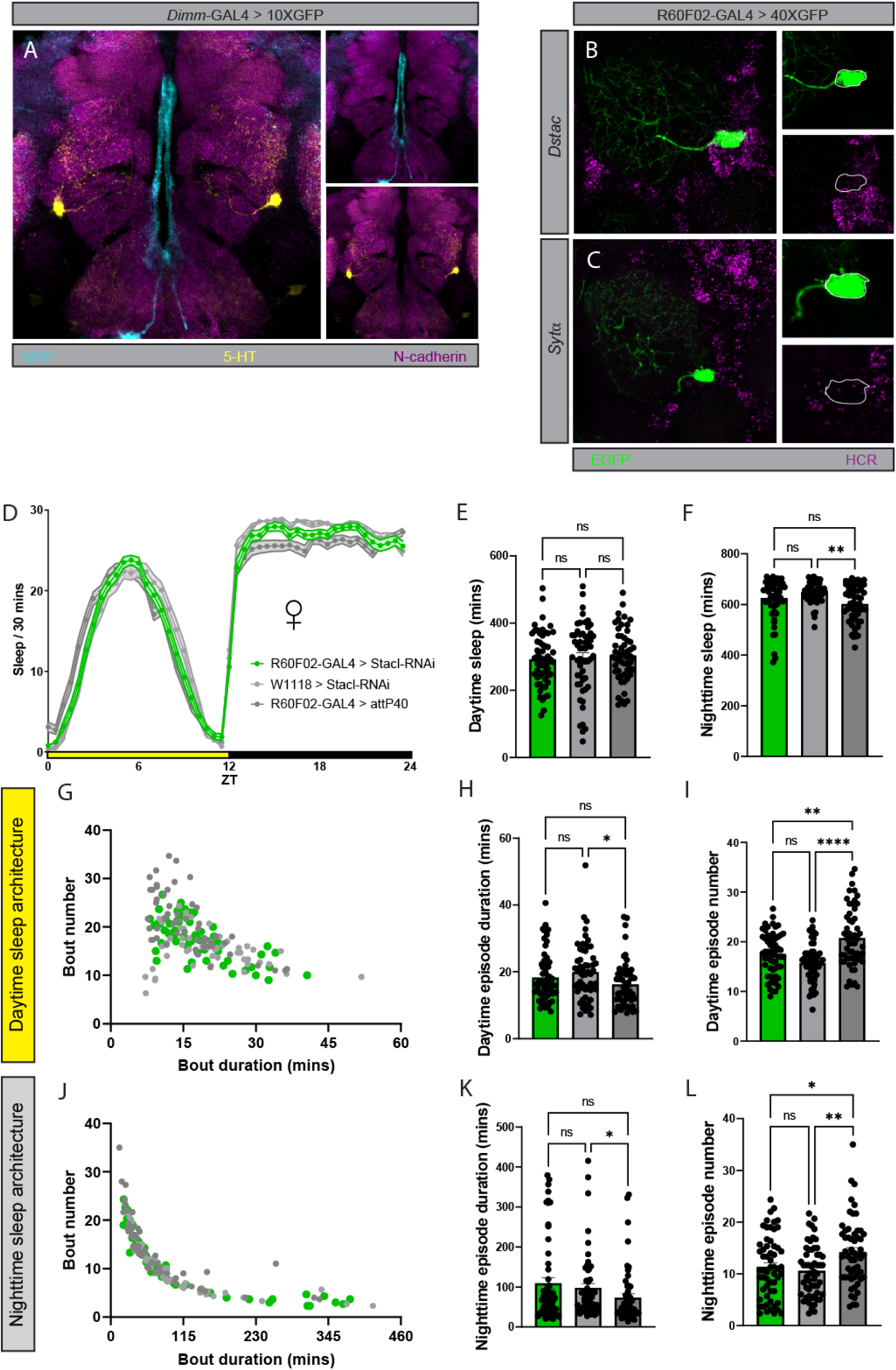
CSDns may express the neuropeptide-related gene *stacl*, but knockdown of *stacl* in the CSDns does not modulate sleep architecture. A) *dimm*-GAL4 expression pattern while co-labeling for 5-HT. GFP signal is driven in the GAL4 (cyan), 5-HT (yellow), neuropil labeled by N-cadherin (magenta). **B-C**) *stacl* and *syt*-ɑ mRNA labeling in the CSDns while co-labeling R60F02-GAL4 eGFP. **D**) Sleep per 30 minutes of flies where CSDn *stacl* expression is reduced via Dicer, R60F02-GAL4 > *stacl*-RNAi. Yellow bar represents daytime (ZT 0-12), while black is nighttime (ZT 12-24). **E-F**) Average sleep duration of **E**) daytime and **F**) nighttime sleep phases. **G**) Scatter plot of the per-fly daytime sleep architecture across each tested genotype. **H**) Average daytime sleep bout duration. **I**) Average daytime sleep bout number. **J**) Scatter plot of the per-fly nighttime sleep architecture across each genotype. **K**) Average nighttime sleep bout duration. **L**) Average nighttime sleep bout number. In each behavior experiment, 3-7 day age mated male flies were tested, N=55-58 per genotype. Lightest gray = Dicer and *stacl*-RNAi control, darker gray = Dicer, R60F02-GAL4 control, green = Dicer, R60F02-GAL4 > *stacl*-RNAi. See methods for details on statistical analysis. *p<0.05, **p<0.01, ***p<0.001, ****p<0.0001.

